# Nuclear reassembly defects after mitosis trigger apoptotic and p53-dependent safeguard mechanisms in *Drosophila*

**DOI:** 10.1101/2024.01.21.576567

**Authors:** Jingjing Li, Laia Jordana, Haytham Mehsen, Xinyue Wang, Vincent Archambault

## Abstract

In animals, mitosis involves the breakdown of the nuclear envelope and the sorting of individualized, condensed chromosomes. During mitotic exit, emerging nuclei reassemble a nuclear envelope around a single mass of interconnecting chromosomes. The molecular mechanisms of nuclear reassembly are incompletely understood. Moreover, the cellular and physiological consequences of defects in this process are largely unexplored. Here, we have characterized a mechanism essential for nuclear reassembly in *Drosophila*. We show that Ankle2 promotes the PP2A-dependent recruitment of BAF and Lamin at reassembling nuclei, and that failures in this mechanism result in severe nuclear defects after mitosis. We then took advantage of perturbations in this mechanism to investigate the physiological responses to nuclear reassembly defects during tissue development *in vivo*. Partial depletion of Ankle2, BAF or Lamin in imaginal wing discs results in wing development defects accompanied by apoptosis. We found that blocking apoptosis strongly enhances developmental defects. Blocking p53 does not prevent apoptosis but enhances defects due to the loss of a cell cycle checkpoint. Our results suggest that apoptotic and p53-dependent responses play a crucial role in safeguarding tissue development in response to sporadic nuclear reassembly defects.

## INTRODUCTION

The passage through mitosis requires a profound transformation of the nucleus. In many eukaryotes including humans, the nuclear envelope (NE) is broken, allowing cytoplasmic centrosomes and chromosomes to connect through a bipolar spindle of microtubules (1, 2). In parallel, replicated chromosomes become condensed and individualized, allowing their correct bipolar attachment and the segregation of sister chromatids on the spindle. Nuclear envelopes and the associated lamina then reassemble around the two segregated groups of chromosomes that decondense, generating daughter nuclei (3, 4). The reassembly of a nucleus after mitosis is essential for cell viability, cell proliferation and development. However, errors in nuclear reassembly (NR) can lead to partial structural defects of the nucleus that may include aberrantly shaped nuclei, micronuclei and an abnormal lamina. Nuclei harboring such flaws may undergo abnormal nucleocytoplasmic exchanges, abnormal gene expression, DNA damage or disintegration (5, 6). However, how cells and tissues respond to NR defects *in vivo* is still unclear.

The lamina is a protein network at the interface between chromatin and the NE in animal cells (7). It confers structure and rigidity to the nucleus, and it contributes to organize chromatin domains and regulate transcription (8). The lamina can be composed of A/C-type lamins and B-type lamins. These proteins form dimers that further assemble into higher-order structures (7, 9). Mutations in lamin genes cause congenital diseases referred to as laminopathies that include various forms of progeria and dystrophies (9, 10). At the cellular level, these mutations often cause an irregular lamina and an abnormal nuclear shape. Lamina defects are associated with a propensity of the NE to form blebs and to rupture (11). Lamina down-regulation and abnormalities are also associated with normal aging and with cancer (12–14).

Micronuclei frequently occur in cancer cells, where their NE often breaks (15, 16). Micronuclei compromise the stability of the chromosomes they contain as they are the site of extensive DNA damage (17). When they break, their DNA can activate a cGAS-STING-dependent innate immune response in mammals (18). Other studies suggested that micronuclei can alternatively trigger apoptosis (5, 19). Micronuclei were also linked to cellular senescence (20). Some nuclear defects can be repaired. A ruptured NE can be resealed by the ESCRT-III membrane fusion machinery (6). Micronuclei can reintegrate the main nucleus (5). The importance of the various responses to NR defects in the context of tissue development *in vivo* is unclear. Investigating this question requires the ability to experimentally interfere with NR mechanisms specifically in a developing animal.

In recent years, the molecular mechanisms that govern NR after mitosis have become better understood. A central pathway in this process involves the protein Barrier-to-Autointegration Factor (BAF) (21). This dimeric DNA-binding protein is recruited to chromosomes in telophase and links chromosomes to each other as they initiate the reassembly of a single nucleus (22–26). In addition, BAF interacts with Lamin A/C and with transmembrane proteins of the NE that contain a Lap2-Emerin-Man1 (LEM) domain (27–30). In human cells, the loss of BAF strongly increases their propensity to micronucleation during NR (22). As cells enter mitosis, the phosphorylation of BAF by vaccinia-related kinases (VRKs) induces the dissociation of BAF from DNA (31, 32). During mitotic exit, the dephosphorylation of BAF is required for its recruitment to chromosomes. This dephosphorylation depends on Protein Phosphatase 2A (PP2A) and on its co-factor Ankle2/Lem4 (33, 34). The PP2A-Ankle2-BAF pathway appears to play a conserved role in NR at least in *H. sapiens* (humans), *C. elegans* (round worms) and *D. rerio* (fish) (33, 35). BAF is conserved in *Drosophila melanogaster*, where it is also required for NR (36, 37). In this system, the phosphorylation of BAF by the VRK ortholog Ballchen/NHK-1 is required for its dissociation from DNA in mitosis and female meiosis (37, 38). We have previously shown that BAF dephosphorylation and recruitment to reassembling nuclei depends in part on PP2A with its B55/Tws regulatory subunit (37). An Ankle2 ortholog exists in *Drosophila* and was shown to promote asymmetric cell divisions in the developing larval brain (39). Consistent with this function, hypomorphic mutations *ANKLE2* cause microcephaly in humans (40, 41). However, whether *Drosophila* Ankle2 is required in NR has not been investigated.

Here, we show that Ankle2 promotes BAF recruitment and NR during mitotic exit. Disruption of Ankle2 or BAF leads to defective nuclei with lamina defects and dispersed, abnormally condensed chromatin. We took advantage of these perturbations to examine the cellular and physiological consequences of NR defects in a proliferative tissue *in vivo*. We found that disrupting Ankle2, BAF or Lamin function in larval imaginal wing discs results in defective NR, apoptosis and developmental defects in adult wings. Moreover, we found that disrupting apoptosis or a p53-dependent response in this context strongly enhances tissue development defects. Thus, these responses play essential roles in promoting the development of a proliferative tissue that incurs frequent structural nuclear defects after mitosis.

## RESULTS

### *Drosophila* Ankle2 promotes BAF dephosphorylation and recruitment during nuclear reassembly after mitosis

In *C. elegans* and human cells, Ankle2 promotes BAF recruitment during NR (33). Whether Ankle2 functions in a similar manner in *Drosophila* has not been examined. We used *Drosophila* cells in culture to test if Ankle2 is required for BAF recruitment. RNAi depletion of Ankle2 was verified by Western blot using newly generated custom antibodies (Fig 1A). We then examined the localization of N-terminally Flag-tagged BAF (Flag-BAF) (Fig 1B). In control cells, Flag-BAF was enriched in the nucleus relative to the cytoplasm, consistent with the previously observed localization of endogenous BAF (36). We found that this nuclear enrichment was lost after Ankle2 depletion (Fig 1C). Moreover, while Flag-BAF was enriched at the NE relative to the nucleoplasm in control cells, this enrichment was lost after Ankle2 depletion (Fig 1D). To visualize BAF recruitment during mitotic exit, we used video microscopy with D-Mel cells expressing GFP-BAF and mCherry-Tubulin (Fig 1E and Videos 1-2). During early mitosis, GFP-BAF was dispersed in the cytoplasm, but it became strongly enriched on segregated chromosomes in telophase, before being restricted to the nuclear periphery in interphase. This dynamics is consistent with previous results (37). By contrast, in Ankle2-depleted cells, GFP-BAF never became strongly enriched on chromosomes in telophase and formed aggregates at the nuclear periphery at a later stage.

**Figure 1.**
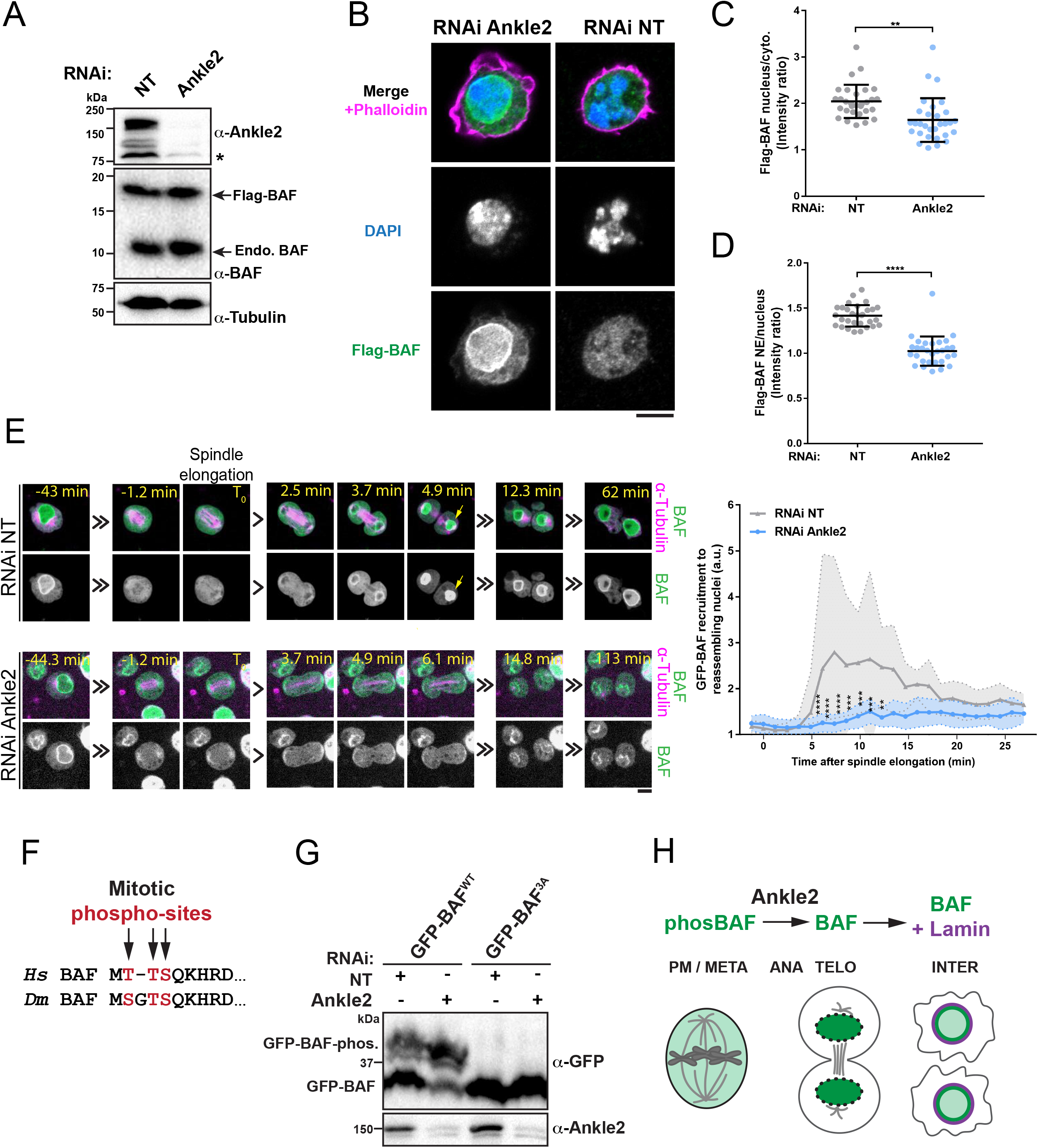
Ankle2 is required for BAF dephosphorylation and recruitment to reassembling nuclei. **A.** Western blots showing the RNAi depletion of Ankle2 in D-Mel cells expressing Flag-BAF. NT: Non-target RNAi. *Non-specific band. The RNAi-sensitive bands likely correspond to Ankle2 splice variants. **B.** Immunofluorescence showing the subcellular localization of Flag-BAF in cells after RNAi depletion of Ankle2 and in control cells (NT RNAi). Cells were also stained with phalloidin to reveal actin. **C.** Quantification of the nucleus/cytoplasm fluorescence intensity ratios in cells after Ankle2 RNAi or NT RNAi. **D.** Quantification of the nuclear envelope/nucleus (nucleoplasm) fluorescence intensity ratios in cells after Ankle2 RNAi or NT RNAi. **E.** Left: Live imaging of mitosis in cells expressing GFP-BAF (green) and mCherry-Tubulin (magenta) after Ankle2 RNAi or NT RNAi. T_0_ was set as the beginning of spindle elongation in anaphase. The strong recruitment of GFP-BAF on DNA in telophase observed in control cells (yellow arrow), does not occur in Ankle2 RNA cells. Right: Quantification of GFP-BAF fluorescence intensity at reassembling nuclei. All scale bars: 5 μm. All error bars: S.D. **p < 0.01, **** p < 0.0001 from unpaired t-tests. **F.** Conservation of the mitotic phosphorylation sites in the N-terminus of BAF in humans (*Hs*) and *Drosophila* (*Dm*). **G.** BAF phosphorylation in its N-terminus depends on Ankle2. Cells expressing GFP-BAF WT or 3A (S2A, T4A, S5A) were treated for Ankle2 RNAi or NT RNAi. Extracts were submitted to SDS-PAGE with the addition of Phos-tag to increase mobility shifts due to phosphorylation. Westerns blots for GFP and Ankle2 are shown. Note that slow-migrating bands corresponding to phosphorylated GFP-BAF (GFP-BAF-phos.) are eliminated by the 3A mutations and are increased upon Ankle2 RNAi. **H.** Ankle2 promotes BAF dephosphorylation and recruitment to reassembling nuclei during mitotic exit, and BAF-dependent recruitment of Lamin.

While BAF phosphorylation at conserved N-terminal sites (Ser2, Thr4 and/or Ser5 in *Drosophila*) promotes its release from DNA in mitosis, BAF recruitment to DNA in telophase requires the dephosphorylation of these sites (Fig 1F) (31, 37, 38). We observed multiple bands for GFP-BAF on Western blot after SDS-PAGE in the presence of Phos-tag (Fig 1G). Mutation of the phosphorylation sites into alanine residues abolished the upper bands, indicating that they correspond to phosphorylated BAF. We found that depletion of Ankle2 led to an increase in the slow-migrating forms of GFP-BAF. Altogether, these results indicate that Ankle2 is required for BAF dephosphorylation and recruitment to reassembling nuclei in *Drosophila* (Fig 1H), a role that is conserved in *C. elegans* and humans (33).

### Ankle2 and BAF promote nuclear reassembly

In human cells, BAF was shown to prevent micronucleation during mitotic exit (22). BAF is also required for Lamin A recruitment (26, 42). To test if BAF plays a similar role in *Drosophila*, we depleted BAF by RNAi in D-Mel cells and examined nuclear morphology after staining DNA and Lamin. To maximize the penetrance of the phenotypes, we transfected dsRNA once or twice, analyzing cells after 4 days or 7 days of depletion, respectively (Fig S1A). We found that BAF depletion results in nuclei with abnormal morphology, often with apparent micronuclei and hypercondensed DNA, when assessed from DAPI staining (Fig 2A-C). In addition, Lamin was often mislocalized, forming aggregates (Fig 2A, D). Thus, BAF is required for normal NR, promoting normal nuclear morphology and Lamin recruitment. Since Ankle2 is required for BAF recruitment, and BAF is required for NR, we tested if Ankle2 was also required for NR. We found that RNAi depletion of Ankle2 induced aberrant nuclei similarly to BAF depletion (Fig 2A-D).

**Figure 2.**
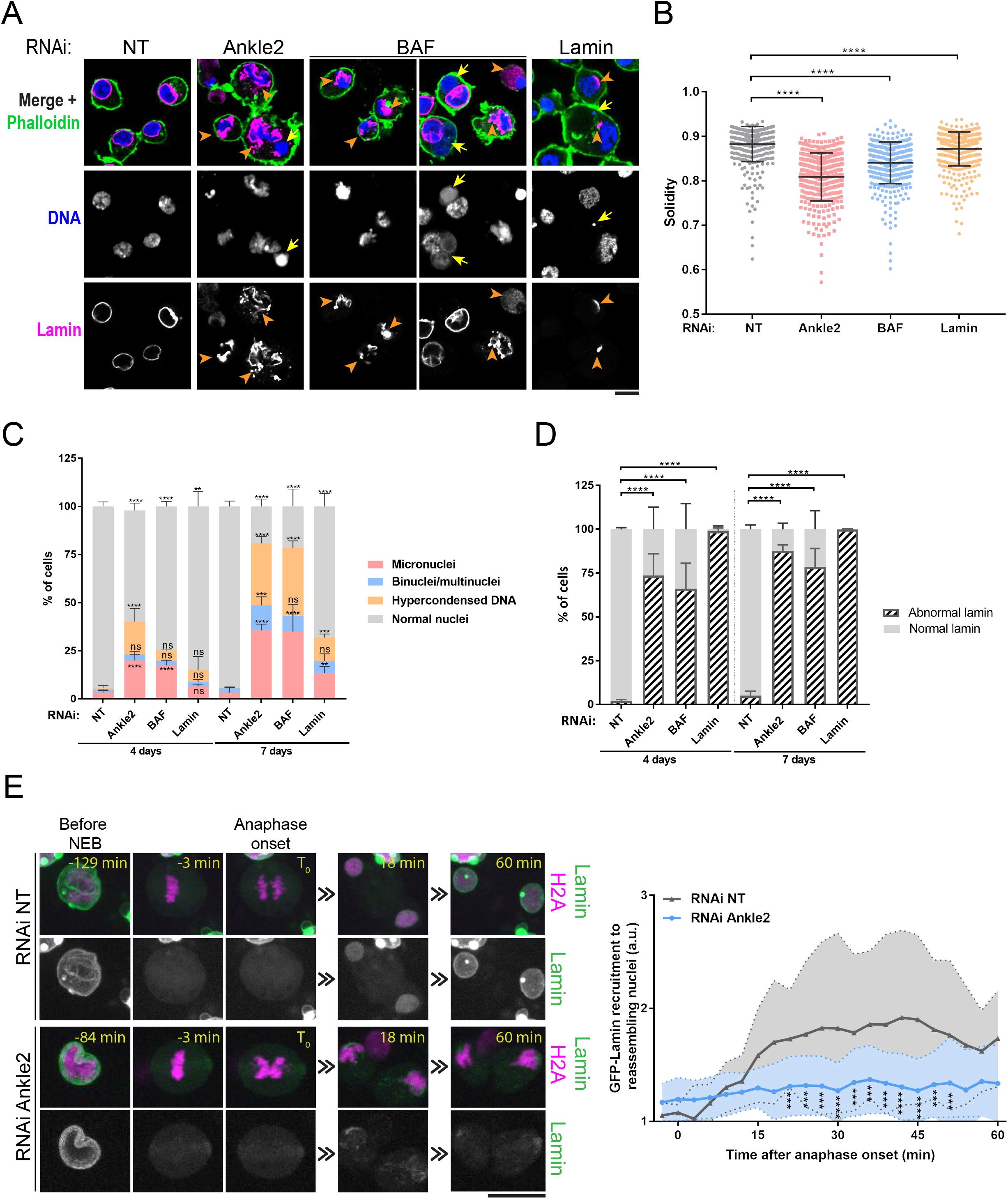
Ankle2, BAF and Lamin are required for nuclear reassembly in cells in culture. **A.** D-Mel cells were treated by RNAi for 4 days to deplete Ankle2, BAF or Lamin. Cells were analyzed by immunofluorescence to reveal Lamin (magenta), DNA (DAPI, blue) and actin (phalloidin, green). NT: Non-Target RNAi. After depletion of Ankle2 or BAF, DNA and Lamin are disorganized. After partial depletion of Lamin, cells appear devoid of Lamin or display small patches of Lamin. Yellow arrows: apparent micronuclei without Lamin. Orange arrowheads: Lamin aggregates or patches. **B.** Quantification of nuclear solidity (area / convex surface) based on the DNA staining from cells treated as in A. **C.** Quantification of nuclear phenotypes based on the DNA staining from cells treated by RNAi as indicated. **D.** Quantification of Lamin phenotypes from cells treated by RNAi as indicated. **E.** Left: Live imaging of mitosis in cells expressing Lamin-GFP (green) and H2A-RFP (magenta) after Ankle2 RNAi or NT RNAi. T_0_ was set as the time of anaphase onset. Right: Quantification of GFP-Lamin fluorescence intensity at reassembling nuclei. All scale bars: 5 μm. All error bars: S.D. **p < 0.01, *** p < 0.001, **** p < 0.0001, ns: non-significant from unpaired t-tests.

To visualize if nuclear defects due to Ankle2 depletion arise during mitosis, we used video microscopy with D-Mel cells expressing H2A-RFP and GFP-Lamin. We found that Ankle2 depletion caused an increase in micronucleation occurring at the end of mitosis (Fig S1B-C). We previously showed that Lamin recruitment during NR depends on BAF recruitment (37). Consistent with this notion, we found that Ankle2 depletion led to reduced GFP-Lamin recruitment after mitosis (Fig 2E, Videos 3-4 and Fig S1B-C). However, depleting Lamin itself caused much less nuclear morphology defects than depleting Ankle2 or BAF (Fig 2A-D). This is consistent with a Lamin-independent role of BAF in cross-bridging chromosomes during telophase to promote the normal morphology of nascent nuclei (22). We conclude that Ankle2 is required for the recruitment of BAF and Lamin (Fig 1H), and for normal NR after mitosis in *Drosophila*.

### Depletion of Ankle2 or BAF in wing discs causes nuclear defects and wing development defects

To investigate the cellular and functional consequences of NR defects during tissue development, we used genetic manipulations in imaginal wing discs. These tissues rely on cell proliferation at the larval stage to generate adult wings. To silence the expression of target genes in this tissue, we used the Gal4-UAS system. Nubbin-Gal4 (Nub-Gal4) was used to drive the expression of RNAi constructs under the control of UAS. Nub-Gal4 is expressed specifically in the pouch area of the discs which is destined to develop into the wing proper (43). We found that depletion of Ankle2 causes nuclear defects in the wing pouch. Immunofluorescence revealed nuclei with irregular lamina surrounding DNA, hypercondensed DNA, masses of DNA devoid of Lamin, and Lamin aggregates separated from DNA (Fig 3A). In parallel to those nuclear defects, adult wings that developed under Nub-Gal4-driven depletion of Ankle2 were smaller than control wings, or almost absent, depending on which RNAi line was used (Fig 3B and S2).

**Figure 3.**
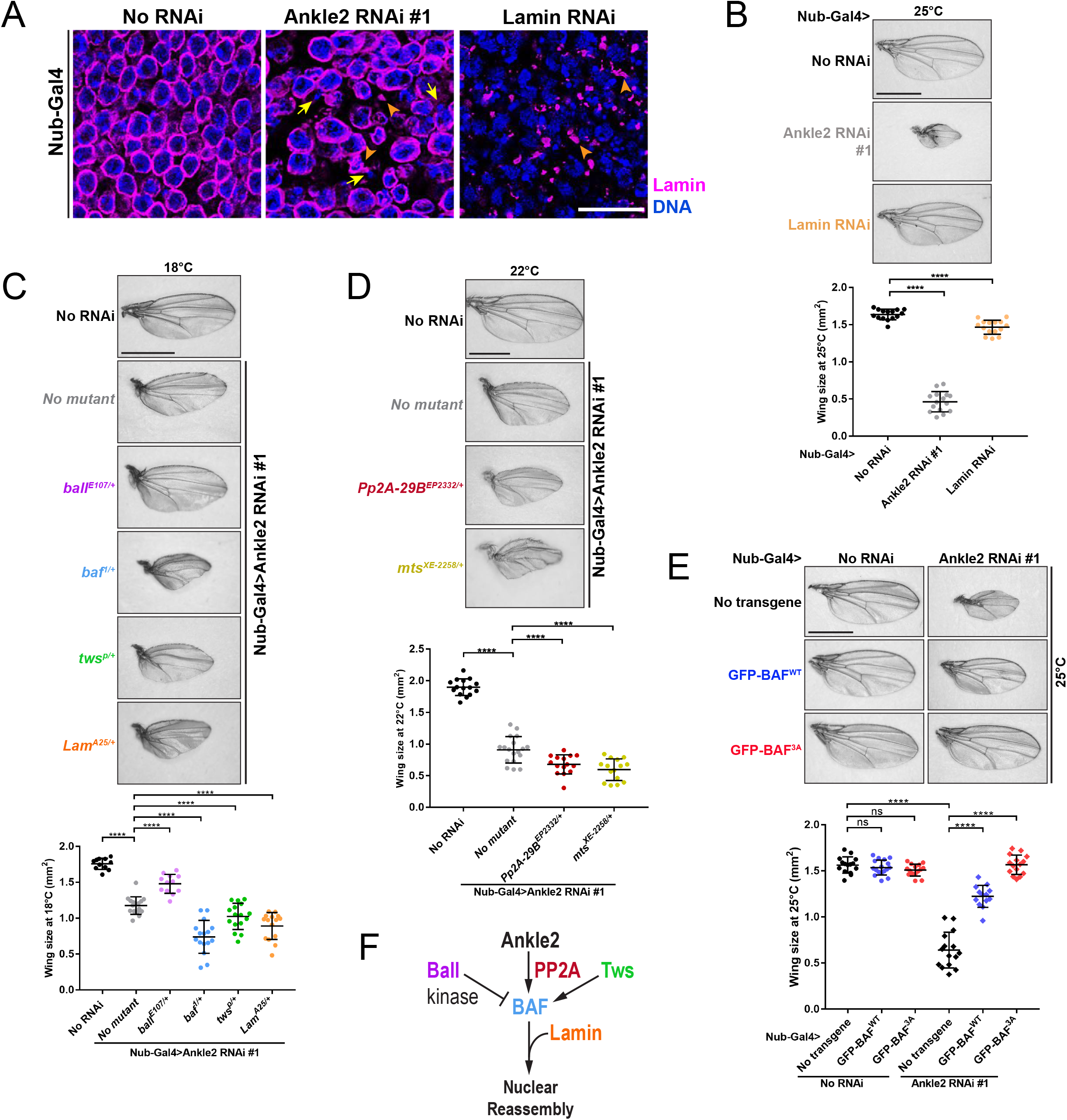
Disruption of Ankle2 function in BAF regulation during wing development causes nuclear defects and adult wing defects. **A.** Nuclear phenotypes in wing discs after induction of Ankle2 RNAi, Lamin RNAi or no RNAi in the pouch area of the wing disc using Nub-Gal4 at 25°C. Wing discs were analyzed by immunofluorescence against Lamin (magenta) and DNA (DAPI, blue). Yellow arrows: DNA without Lamin. Orange arrowheads: Lamin aggregates or patches. Scale bars: 10 μm. **B.** Depletion of Ankle2 or Lamin in wing discs results in smaller wings. Top: Examples of adult wings after induction of Ankle2 RNAi, Lamin RNAi or no RNAi in the pouch area of the wing disc using Nub-Gal4 at 25°C. Bottom: Quantification of wing sizes (surface area). **C.** Mutations in *baf*, *tws* and *lamin* enhance, while a mutation in *ball* suppresses, the small wing phenotype resulting from Ankle2 depletion. Top: Examples of adult wings of the indicated genotypes after induction of Ankle2 RNAi using Nub-Gal4 at 18°C. Bottom: Quantification of wing sizes at 18°C or 22°C. **D.** Mutations in PP2A genes enhance the small wing phenotype resulting from Ankle2 depletion. Top: Examples of adult wings of the indicated genotypes after induction of Ankle2 RNAi using Nub-Gal4 at 22°C. Bottom: Quantification of wing sizes. **E.** Expression of GFP-BAF^3A^ driven by Nub-Gal4 rescues the small wing phenotype due to Ankle2 depletion. Top: Examples of adult wings of the indicated genotypes at 25°C. Bottom: Quantification of adult wing sizes. In all experiments, Ankle2 depletion was done using line VDRC100655. All scale bars for adult wings: 1 mm. All error bars: S.D. **** p < 0.0001, ns: non-significant from unpaired t-tests with Welch’s correction. **F.** Conceptual representation of the molecular network controlling NR (see text for details).

Depletion of BAF using two RNAi lines yielded nuclear defects similar to those obtained after Ankle2 RNAi (Fig S3A). The line that resulted in the strongest defects (VDRC102013) has its UAS RNAi construction inserted at cytolocation 40D that is known to also induce Gal4-dependent overexpression of the transcription factor Tio, which can result in development defects (44). However, a control line with a UAS insertion at the same site (leading to Tio overexpression only; VDRC60101) did not result in similar nuclear defects (Fig S3A). Thus, nuclear defects observed upon Nub-Gal4-driven depletion of BAF are specifically due to BAF depletion and not Tio overexpression. However, both the BAF RNAi 40D insertion an its Tio-overexpressing control insertion resulted in very small adult wings (Fig S3B-C). We conclude that this adult wing phenotype is not specific to BAF depletion for this RNAi line. Nevertheless, depletion of BAF with an alternative RNAi construction (line BDSC36018) resulted in both nuclear defects and reduced adult wing size (Fig S3A-C).

Partial depletion of Lamin in wing discs resulted in nuclei lacking Lamin around a large fraction of their periphery, with the remaining Lamin forming foci, similarly to the phenotype observed in cell culture (Fig 3A). This depletion of Lamin also resulted in a reduction of adult wing size, although to a lesser extent than Ankle2 depletion (Fig 3B). Depletion of Ankle2, BAF or Lamin also caused delocalization of Nuclear Pore Complex (NPC) proteins and Otefin (an inner nuclear membrane protein with a LEM domain that interacts with BAF) from the NE (Fig S3D). Therefore, depletion of Ankle2, BAF or Lamin in the developing wing causes nuclear structure defects associated with a reduction in adult wing size.

### Genetic validation of an Ankle2-BAF-Lamin pathway required for nuclear reassembly

To identify functional interactions of the Ankle2-BAF-Lamin pathway, we exploited the small wing phenotype. We used depletion of Ankle2 with the weaker RNAi line (VDRC100655) which resulted in a partial phenotype due to incomplete penetrance, allowing the identification of enhancer and suppressor genes. To facilitate the identification of enhancer and suppressor genes, we could also modulate the phenotype by varying the temperature because Gal4-UAS-dependent expression increases with temperature. As expected, wing size was further reduced by the introduction of one mutant allele of *baf*, consistent with the role of Ankle2 in promoting BAF recruitment during NR (Fig 3C). Conversely, wing size was partially rescued by introduction of one mutant allele of *ball*/*nhk-1*, which encodes the kinase that phosphorylates BAF to induce its dissociation from DNA in M-phase (38). Mutant alleles of *mts* and of *Pp2A-29B*, which encode the catalytic and structural subunits of PP2A, respectively, enhanced the small wing phenotype (Fig 3D). These results are consistent with the role of PP2A in dephosphorylating BAF to promote its recruitment during NR, and with the interaction of PP2A with Ankle2 in this process. The small wing phenotype was also enhanced by one mutant allele of *tws*, which encodes the B55 regulatory subunit of PP2A (Fig 3C). This is consistent with our previous finding of a role of PP2A-Tws in the dephosphorylation and recruitment of BAF during NR (37). Heterozygosity for mutations in *lamin* also enhanced the phenotype (Fig 3C).

To test if wing development defects upon Ankle2 depletion are due to a failure to dephosphorylate BAF, we overexpressed non-phosphorylatable GFP-BAF^3A^. We found that Nub-Gal4-dependent expression GFP-BAF^3A^ in the wing pouch fully rescued wing size when Ankle2 is depleted (Fig 3E). By contrast, expression of GFP-BAF^WT^ rescued wing size only partially. Altogether, these results support the notion that Ankle2 functions with PP2A in promoting the dephosphorylation and recruitment of BAF during NR and that a failure in this function results in developmental defects *in vivo*. Overall, the genetic interactions we observed in the context of wing development are completely consistent with an emerging molecular network where Ankle2 plays a conserved role in NR (Fig 3F).

### Nuclear reassembly defects trigger apoptosis

The nuclear defects observed upon depletion of Ankle2, BAF or Lamin (fragmented nuclei, hypercondensed DNA, abnormal Lamin, Fig 2 and 3A) are reminiscent of pyknotic nuclei that develop during apoptosis. We therefore sought to determine whether depletion of Ankle2, BAF or Lamin induces apoptosis. The effector Caspase 3, called Dcp-1 in *Drosophila*, is activated downstream of the canonical apoptosis pathway (Fig 4A) (45, 46). Staining for cleaved, activated Dcp-1 (hereafter Dcp-1) revealed a significant increase in apoptosis after depletion of Ankle2, BAF or Lamin (Fig 4B-C and S4A-B). The intensity of Dcp-1 signals mirrored the penetrance of the small-wing phenotype with two Ankle2 RNAi lines (Fig S2 and S5A-C). An increase in apoptosis was also observed after depletion Otefin or Man1, two structural LEM-D proteins of the NE that interact with BAF (Fig 4B-C). Moreover, the Dcp-1 signal tended to be located in areas with defective nuclei (Fig S5B). Increased apoptosis was also defected by TUNEL assay in wing discs depleted of Ankle2, BAF or Lamin (Fig 4D-F, S4C-D). TUNEL and Dcp-1 signals tended to increase together, and foci were often observed in close proximity suggesting that they tend to occur in the same cells (Fig 4E-F). Depletion of Ankle2 or BAF in D-Mel cells in culture also increased Dcp-1 signals concomitantly with hypercondensed DNA in pyknotic nuclei (Fig S6). Altogether, these results indicate that disruption of NR triggers apoptosis.

**Figure 4.**
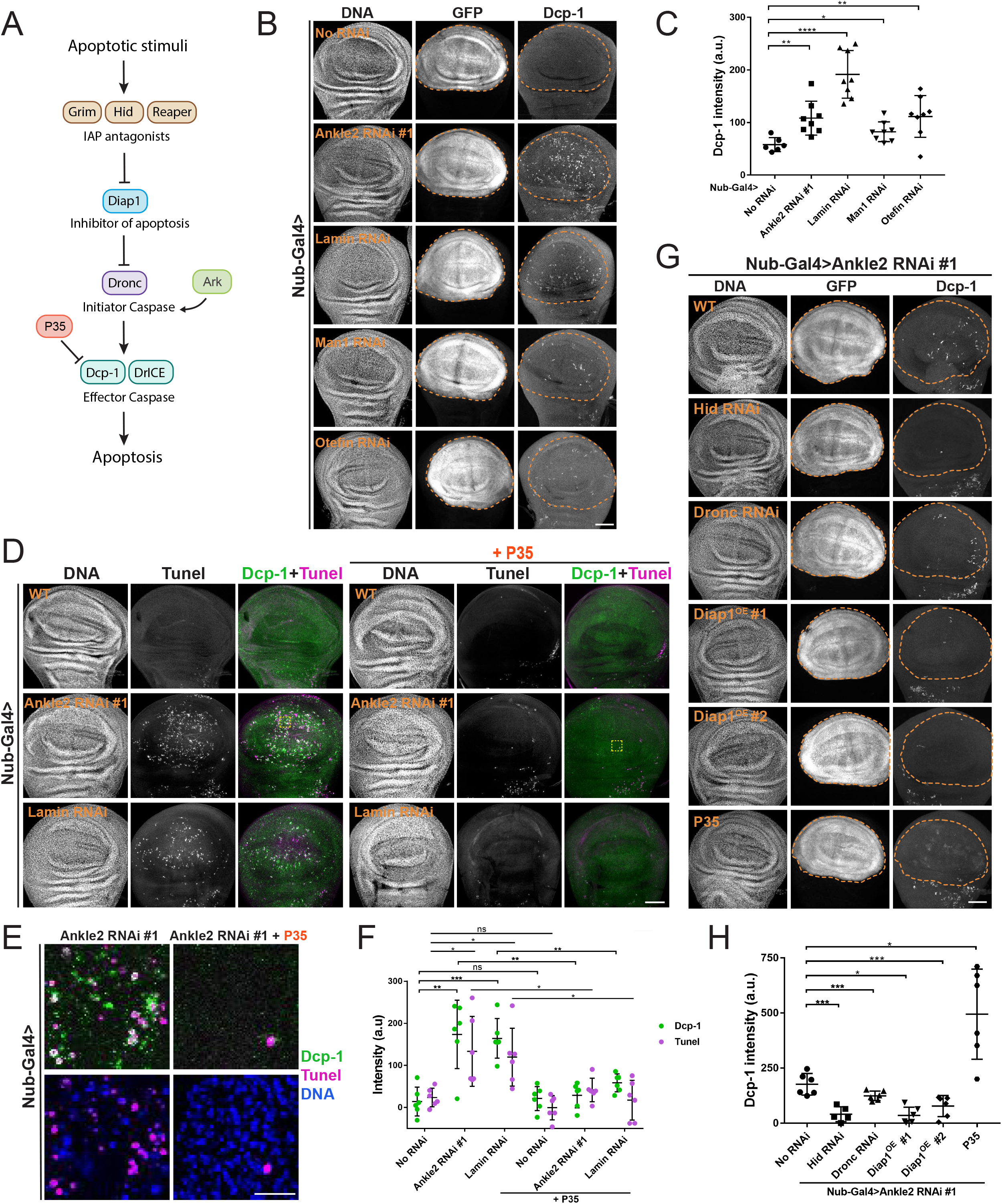
Perturbation of nuclear reassembly mechanisms induce apoptosis. **A.** The classical apoptotic pathway in *Drosophila*. **B.** Depletion of Ankle2, Lamin or BAF interactors of the nuclear envelope induces apoptosis. RNAi for the indicated proteins was induced in the wing pouch by Nub-Gal4 at 25°C. In parallel, UAS-GFP was used as a marker of the region of interest (wing pouch, inside dotted lines). Wing discs were analyzed by immunofluorescence against cleaved Dcp-1 and stained for DNA (DAPI). **C.** Quantification of Dcp-1 intensities measured in the wing pouch region of individual wing discs of the indicated genotypes as in B. **D.** Detection of apoptosis by TUNEL (magenta) and simultaneous immunofluorescence for cleaved Dcp-1 (green) in wing discs depleted of Ankle2 or Lamin. Expression of P35 (right) abrogates apoptosis. Yellow boxes: Areas enlarged in panel E. **E.** Enlargements of regions of interest from images in D (yellow boxes) Scale bars: 10 μm. **F.** Quantifications of TUNEL and Dcp-1 signals in wing discs of the indicated genotypes. **G.** Genetic perturbations abrogating apoptosis decrease Dcp-1 signals in the Ankle2-depleted wing pouch region (GFP positive, inside dotted lines). **H.** Quantification of Dcp-1 intensities measured in the wing pouch of individual wing discs of the indicated genotypes as in G. Note that a diffuse Dcp-1 signal increases following expression of P35 in Ankle2-depleted wing discs. In all experiments, Ankle2 depletion was done using line VDRC100655. All scale bars for imaginal wing discs: 50 μm. All error bars: S.D. *p<0.05, **p < 0.01, *** p < 0.001, **** p < 0.0001, ns: non-significant from unpaired t-tests with Welch’s correction.

To test whether apoptosis was caspase-dependent, we used P35, a protein known to prevent apoptosis in *Drosophila* by inhibiting effector caspases (47, 48). Expression of P35 in wing discs depleted of Ankle2, BAF or Lamin blocked apoptosis measured by TUNEL, as expected (Fig 4D-F). Interestingly, while expression of P35 eliminated bright Dcp-1 foci, it actually increased a diffuse Dcp-1 signal (Fig 4G-H). This is consistent with the fact that while P35 inhibits effector caspases including Dcp-1, it does not block the initiator caspase Dronc, which cleaves Dcp-1 (49, 50). Persistent, diffuse Dcp-1 signals were previously observed following P35 expression (51, 52). In this context, cells positive for Dcp-1 likely accumulate as they do not complete apoptosis. This increase in diffuse Dcp-1 signal was not detected when this staining was done in combination with TUNEL, for unknown reason (Fig 4F). As expected, interfering with apoptosis upstream of P35 abrogated Dcp-1 signals; RNAi depletion of Dronc or Hid (two activators of apoptosis) or overexpression of Diap1 (inhibitor of apoptosis), decreased Dcp-1 signals in Ankle2-depleted wing discs (Fig 4G-H).

To further validate that the apoptosis observed upon Ankle2 depletion reflects a failure in the Ankle2-BAF pathway, we tested genetic perturbations predicted to enhance or rescue apoptosis. As expected, a mutant allele of *baf* increased Dcp-1 signals, while a mutant allele of *ball* decreased them (Fig S7A). In addition, overexpression of GFP-BAF^WT^ or GFP-BAF^3A^ decreased Dcp-1 signals (Fig S7B).

The occurrence of apoptosis in wing discs depleted of NR factors raised the possibility that the nuclear defects observed were a consequence (downstream), rather than a cause (upstream) of apoptosis. To test this possibility, we examined cells in wing discs where apoptosis was blocked by the expression of P35. Interestingly, P35 did not prevent the occurrence of defective nuclei in wing discs depleted of Ankle2 or BAF (Fig 5A). In fact, P35 expression appeared to enhance nuclear defects. We conclude that these defective nuclei are not a consequence of apoptosis. Instead, defective NR appears to be the trigger of an apoptotic response that allows the elimination of defective cells during tissue development.

**Figure 5.**
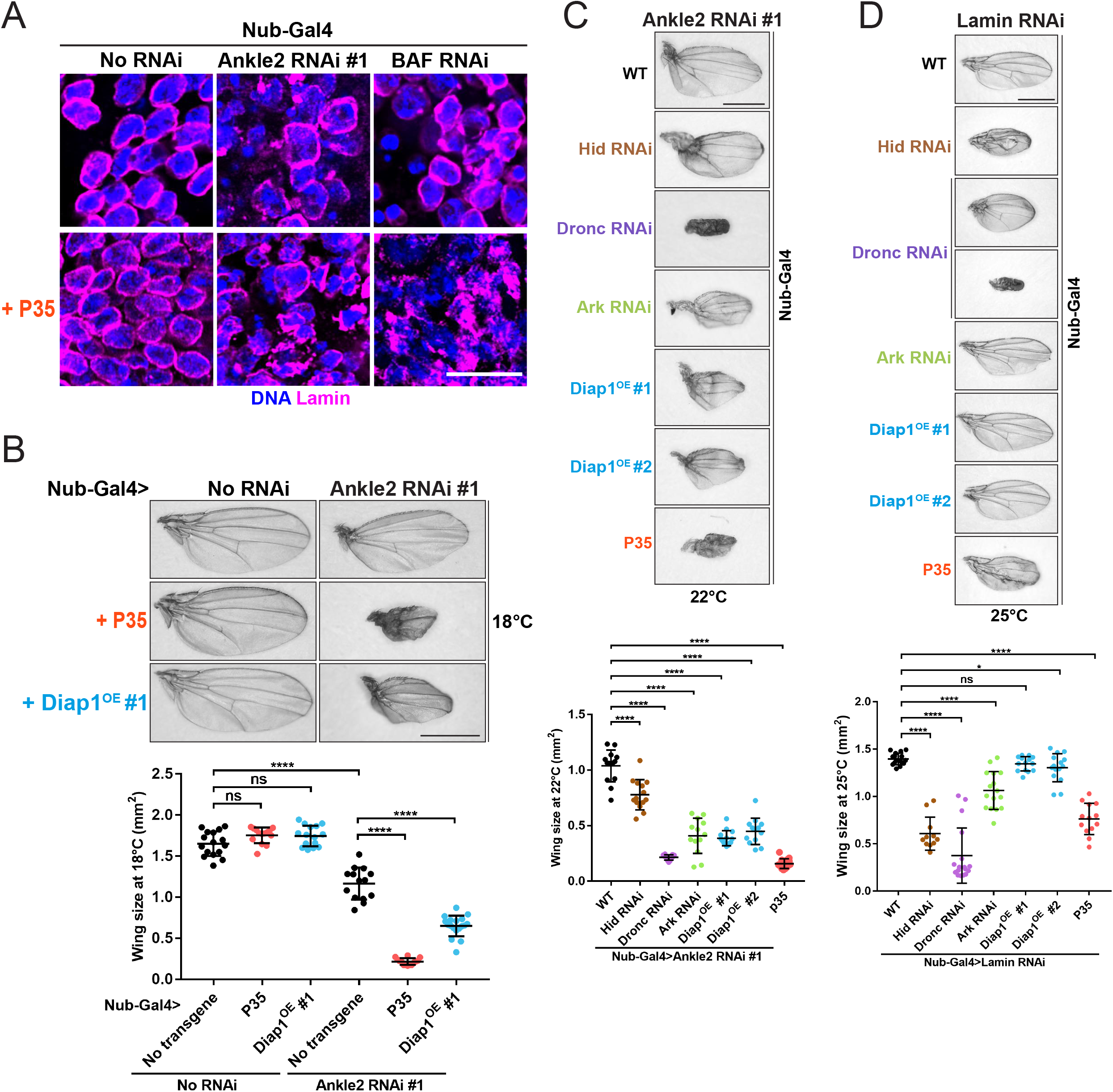
Apoptosis following nuclear reassembly defects promotes normal tissue development. **A.** Expression of P35 in the wing pouch does not eliminate nuclear defects induced by depletion of Ankle2 or BAF. RNAi constructions and P35 expression were driven by Nub-Gal4 at 25°C. Scale bars: 10 μm. **B.** Expression of P35 or overexpression of Diap1 enhances the small wing phenotype resulting from Ankle2 depletion. Top: Examples of adult wings of the indicated genotypes at 22°C. Bottom: Quantifications of wing size from flies of the indicated genotypes at different temperatures. **C-D.** Depletion of positive regulators of apoptosis (Hid, Dronc, Ark) or expression of negative regulators of apoptosis (Diap1, P35), enhances the small wing phenotype resulting from depletion or Ankle2 (C) or Lamin (D). Top: Examples of adult wings of the indicated genotypes. Bottom: Quantifications of wing size from flies of the indicated genotypes. In all experiments, Ankle2 depletion was done using line VDRC100655. All scale bars for adult wings: 1 mm. All error bars: S.D. *p < 0.05, **** p < 0.0001, ns: non-significant from unpaired t-tests with Welch’s correction.

### Apoptosis promotes normal tissue development when nuclear reassembly is compromised

Because apoptosis kills cells, we anticipated that blocking apoptosis may partially rescue the small wing phenotypes resulting from depletion of Ankle2, BAF or Lamin. Strikingly, we obtained the opposite result. Expression of P35 in Ankle2-depleted wing discs enhanced the phenotype, causing adult wings to be even smaller, while expression of P35 alone had no effect on wing size (Fig 5B-C). Moreover, overexpressing Diap1 (negative regulator of apoptosis) or RNAi depletion of Hid, Ark or Dronc (positive regulators of apoptosis) enhanced the small wing phenotype resulting from Ankle2 depletion. Similar results were obtained in the context of the small wing phenotype upon Lamin RNAi (Fig 5D). We conclude that apoptosis promotes normal development when NR defects occur in the wing disc, a proliferative tissue.

### Requirements for apoptosis in response to nuclear reassembly defects

We sought to determine how NR defects trigger apoptosis. NR defects including micronuclei and lamina defects make the NE rupture-prone and can lead to DNA damage (20, 53). Thus, we hypothesized that NR defects may lead to DNA damage that could trigger apoptosis. We found that depletion of Ankle2 in wing discs led to an increase in H2Av phosphorylated at Ser137 (pH2Av), a marker of double-stranded breaks (Fig 6A). It is known that DNA fragmentation can occur as part of the apoptotic process (54). However, blocking apoptosis by expression of P35 did not eliminate the pH2Av staining (Fig 6B). These results suggest that DNA damage occurs as a result of NR defects, upstream of apoptosis.

**Figure 6.**
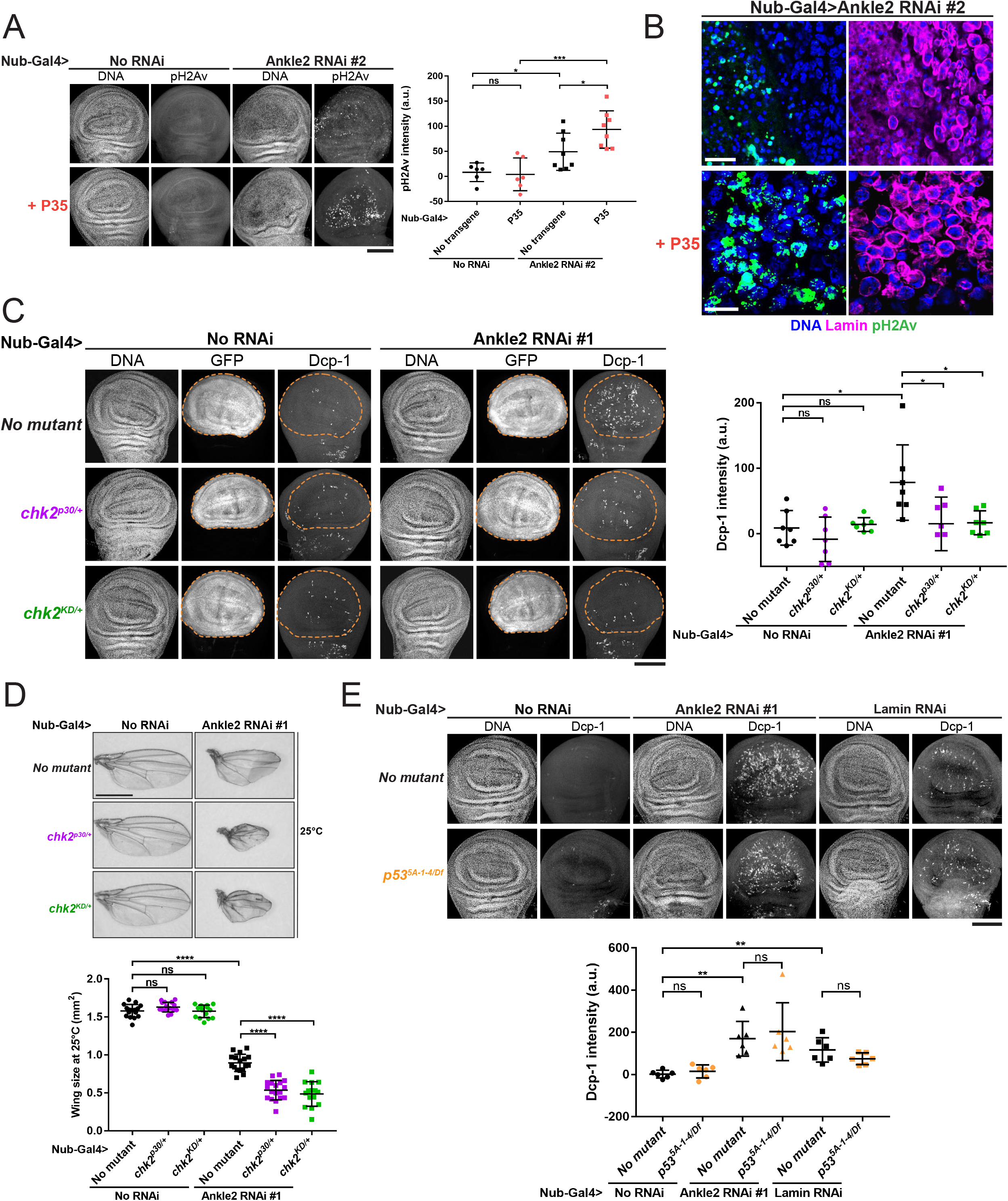
Apoptosis following nuclear reassembly defects occurs downstream of DNA damage and requires Chk2 but not p53. **A.** Depletion of Ankle2 in the wing pouch (line BDSC77437) induces DNA damage independently from apoptosis. Wings discs of the indicated genotypes were stained for pH2Av (pSer137), a marker of DNA double-stranded breaks. Left: Examples of wing discs of indicated genotypes. Right: Quantification of the signal in the entire wing discs. Ankle2 RNAi and P35 expression were driven by Nub-Gal4. **B.** Images of a higher magnification showing pH2Av associated with DNA in defective nuclei after Ankle2 depletion in the wing pouch. Scale bars: 10 μm. **C.** Chk2 is required for apoptosis following Ankle2 depletion in the wing pouch. Left: Examples of wing discs of the indicated genotypes. Right: Quantifications of Dcp-1 signals in the wing pouch (GFP-positive, inside dotted line in images). **D.** Mutations in Chk2 enhance the small wing phenotype resulting from Ankle2 depletion. Top: Example of adult wings of indicated genotype. Scale bar: 1 mm. Bottom: Quantifications of wing sizes at 25°C. **E.** p53 is not required for apoptosis following Ankle2 or Lamin depletion in the wing pouch. Top: Examples of wing discs of the indicated genotypes. Bottom: Quantifications of Dcp-1 signals in the wing pouch (GFP-positive, inside dotted line in images). In experiments of panels C-E, Ankle2 depletion was done using line VDRC100655. All scale bars for whole imaginal wing discs: 50 μm. All error bars: S.D. *<0.05, **p < 0.01, *** p < 0.001, **** p < 0.0001, ns: non-significant from unpaired t-tests with Welch’s correction.

It was previously shown that nuclear lamina defects due to the loss of BAF or Otefin in *Drosophila* female germline stem cells trigger their death in a manner that depends on the DNA damage response kinases ATR and Chk2 (55, 56). An increase in Dcp-1 signal was observed in female germ cells of an *otefin* mutant or upon RNAi depletion of BAF (56). Mutation of *chk2* abolished Dcp-1 signals in *otefin* mutant and decreased Dcp-1 signals in BAF-depleted germ cells (56). We were unable to generate *chk2* null flies with the depletion of Ankle2 in wing discs. However, we found that a single mutant allele of *chk2* was sufficient to decrease Dcp-1 signals (Fig 6C). This is consistent with the previous observation that *chk2* can be haploinsufficient (57). Moreover, heterozygosity for *chk2* mutations enhanced the small-wing phenotype resulting from Ankle2 depletion (Fig 6D). This is consistent with our finding that blocking apoptosis enhanced this phenotype. Thus, our results suggest that Chk2 contributes to the induction of apoptosis in response to NR defects.

In many contexts, the tumor suppressor p53 promotes apoptosis (58, 59). Moreover, p53 is commonly activated by Chk2 for this function in both flies and vertebrates (60–62). We therefore tested if p53 is required for apoptosis induced by NR defects. Surprisingly, we found that inactivation of p53 with mutant alleles or RNAi did not block apoptosis in wing discs depleted of Ankle2 or Lamin (Fig 6E and S8). Altogether our results suggest that NR induces DNA damage and Chk2-dependent apoptosis independently from p53.

### A p53-dependent response promotes tissue development when nuclear reassembly is compromised

Although inactivation of p53 did not eliminate apoptosis upon NR defects, we found that it resulted in an enhancement of the small wing phenotype upon depletion of Ankle2 or Lamin (Fig 7A and S9A). In addition to its role in promoting apoptosis, p53 also functions in arresting the cell cycle in response to various cellular defects and insults (59, 63, 64). In *Drosophila*, p53 is known to arrest the cell cycle by promoting the ubiquitination and degradation of Cyclin E (CycE) (65, 66). To test if this pathway was at play, we simultaneously depleted CycE and Ankle2 by RNAi in developing wings. Strikingly, depletion of both CycE and Ankle2 resulted in larger wings compared to depletion of Ankle2 alone (Fig 7B). Non-target RNAi transgenes were used as controls to exclude the possibility of a rescue resulting from the dilution of Gal4 between two UAS elements. In addition, we found that introduction of one mutant allele of *cycE* also partially rescued wing size upon Ankle2 RNAi (Fig S9B). These results suggest that p53 and CycE function against each other in the context of defects in NR.

**Figure 7.**
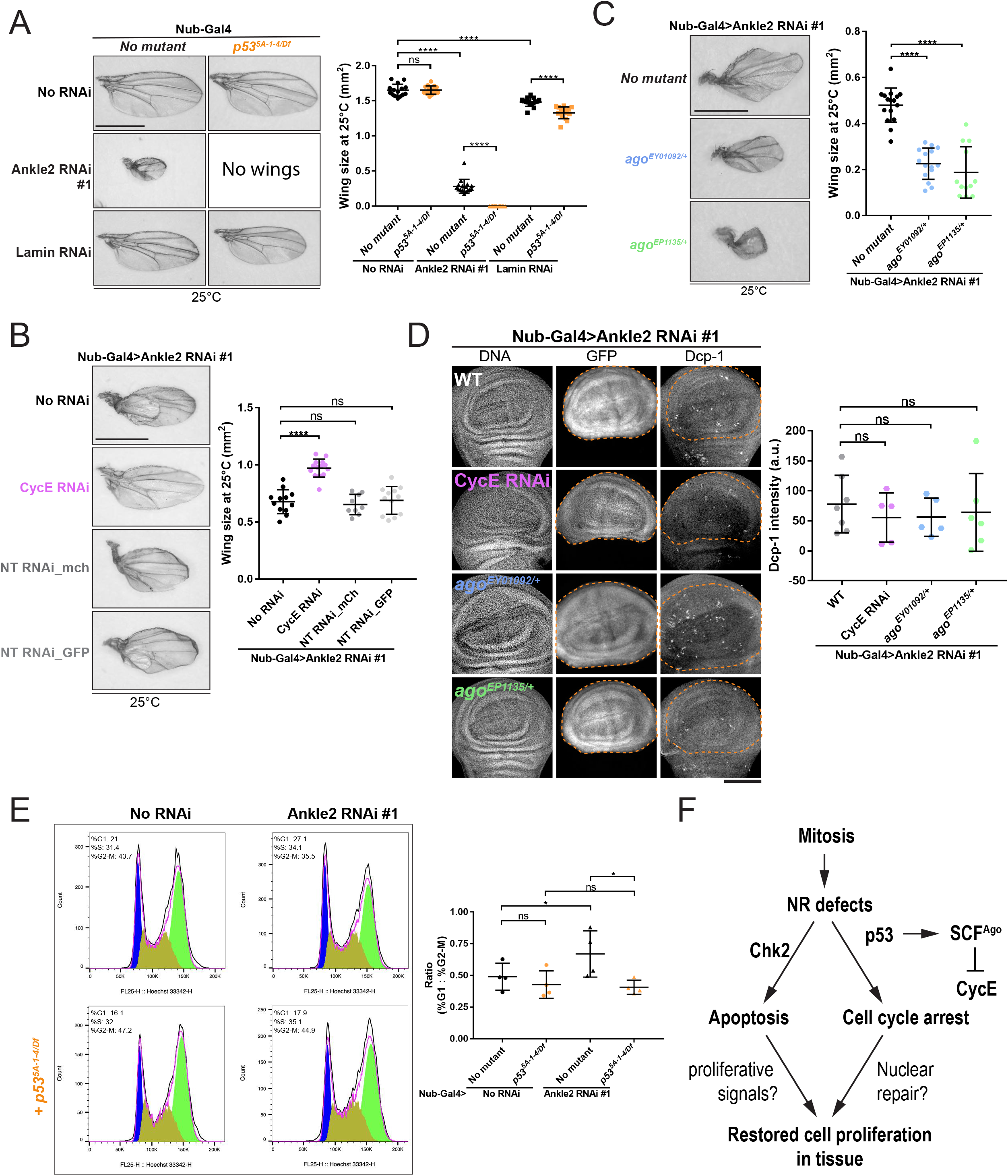
A p53-dependent response promotes tissue development when nuclear reassembly is compromised. **A.** Mutation of *p53* enhances the small wing phenotype resulting from Ankle2 or Lamin depletion. Left: examples of adult wings of the indicated genotypes. Right: quantification of wing sizes. **B.** Simultaneous depletion of CycE rescues the small wing phenotype resulting from Ankle2 depletion. Left: Examples of adult wings of the indicated genotypes. Right: Quantification of wing sizes at 25°C. Non-Target (NT) RNAi constructions against the mCherry (mch) or GFP sequences were used as controls. **C.** Mutations in *Ago* enhance the small wing phenotype resulting from Ankle2 depletion. Left: Examples of adult wings of the indicated genotypes. Right: Quantification of wing sizes at 25°C. **D.** Depletion of CycE or mutation in *Ago* does not modify apoptosis following Ankle2 depletion. Left: Examples of wings discs of the indicated genotypes. Right: Quantification of the Dcp-1 intensities measured in the wing pouch (GFP-positive, inside dotted line). Scale bar: 50 μm. All scale bars for adult wings: 1 mm. All error bars: S.D. **** p < 0.0001, ns: non-significant from unpaired t-tests with Welch’s correction. **E.** Left: Flow cytometry cell cycle analysis (DNA content) of wing disc cells of the pouch area (mCherry positive). Nub-Gal4 was used to drive the depletion of Ankle2 RNAi together with UAS-mCherry, with or without mutation of *p53*. Right: Quantifications ratios of G1/G2-M cells after Nub-Gal4 depletion of Ankle2 with or without mutation of *p53*. *p < 0.05 from paired t-tests. In all experiments, Ankle2 depletion was done using line VDRC100655. **F.** Schematic model showing the p53-dependent cell cycle arrest in response to NR, independently from apoptosis. Both responses may collaborate in restoring normal cell proliferation in a developing tissue.

It was previously shown that p53 promotes CycE degradation by activating expression of Archipelago (Ago), a Fbxw7 family protein that is part of the SCF complex that promotes ubiquitination and subsequent degradation of CycE (65, 66). Consistent with this mechanism being at play in our system, we found that introduction of a single mutant allele of *ago* strongly enhances the small wing phenotype resulting from Ankle2 depletion (Fig 7C). Interestingly, depletion of CycE or mutation of *Ago* in Ankle2-depleted wing discs did not significantly alter apoptotic levels (Fig 7D), consistent with their implication with p53 in a response distinct from apoptosis.

Finally, we tested if NR defects in the wing discs induce an alteration of the cell cycle in a p53-dependent manner. Cell-cycle profile analysis of wing disc cells by flow cytometry revealed that depletion of Ankle2 increases the population of cells in G1 and relative to G2/M (Fig 7E). As expected, mutation of p53 in this context rescued a normal cell cycle profile. Consistent with these observations, a decrease of proliferative cells (EdU+ cells) was also observed in the wing discs upon Ankle2 depletion, and this decreased population of EdU+ cells was also rescued by inactivation of p53 (Fig S9C). These results suggest that in response to NR defects, p53 promotes a cell cycle arrest in G1 via the SCF^Ago^-dependent degradation of Cyclin E, and that this mechanism is independent from apoptosis (Fig 7F).

## DISCUSSION

In this study, we have found that post-mitotic NR in *Drosophila* is mediated by a conserved mechanism that involves Ankle2, PP2A, BAF and Lamin. Using disruptions of this mechanism, we induced NR defects to test the physiological consequences at the cellular and tissue levels. We found that post-mitotic NR defects triggers apoptosis. In addition, we identified a parallel mechanism whereby NR defects trigger a p53-dependent cell cycle arrest. Our results indicate that both responses are beneficial to tissue development.

Previous work in *C. elegans* and human cells has identified Ankle2/Lem4 as key factor for post-mitotic NR (33). Ankle2 interacts with PP2A and promotes the dephosphorylation of BAF that is required for BAF to be recruited to segregated chromosomes and promote NR (33, 34). Recent work on the *Drosophila* ortholog of Ankle2 uncovered its function specifically in the asymmetric divisions of neuroblasts (39). However, whether *Drosophila* Ankle2 functions in NR was not explored. Our results confirm that Ankle2 does function in a conserved manner to promote BAF dephosphorylation and recruitment to reassembling nuclei. Inactivation of Ankle2 or BAF results in post-mitotic NR defects including disorganized DNA and Lamin, which tends to become aggregated and disconnected from DNA. In humans, BAF interacts with Lamin A/C and is required for its recruitment to the lamina (21, 67–70). However, BAF is not required for the recruitment of B-type lamins (70). *Drosophila* Lamin (Dm_0_), the only ubiquitously expressed lamin, is thought to be a closer ortholog to human B-type lamins rather than Lamin A/C (71). Nevertheless, interfering with BAF function in *Drosophila* results in a failure to recruit Lamin, consistent with the fact that BAF forms a complex with Lamin in *Drosophila* (37, 72). However, we found that while depletion of Lamin leads to NR defects, apoptosis and developmental defects, these phenotypes tend to be even stronger when Ankle2 or BAF is depleted. This suggests that the known role of BAF in cross-bridging chromosomes in telophase to ensure the formation of a single nucleus of normal shape (independently of lamins) is a crucial, conserved aspect of its functions in NR (22). In addition, post-mitotic nuclear defects observed after inactivation of BAF could be due in part to its proposed function at centromeres, earlier in mitosis (73, 74).

Altogether, our results suggest that post-mitotic NR defects leads to DNA damage that triggers Chk2-dependent apoptosis. Defective nuclei may harbor holes in their NE or may be more prone to rupture, as demonstrated for micronuclei and nuclei with defective lamina (20, 53). In both cases, DNA would become exposed to cytoplasmic nucleases. The implication of the DNA damage response kinase Chk2 in triggering apoptosis is consistent with previous studies showing that nuclear lamina defects in *Drosophila* female germ cells induce apoptosis through ATR and Chk2 (55, 56). A recent report also showed that a hypomorphic point mutation in BAF (orthologous to a mutation causing progeria in humans) induced Chk2-dependent apoptosis in *Drosophila* wing discs (74). Because Chk2 was shown to induce apoptosis through p53 in other contexts, we expected p53 to also be required for apoptosis following NR defects (60–62). However, we found that p53 is dispensable for this process. Instead, our results suggest that p53 is required for a parallel response where NR-defective cells activate a checkpoint that leads to an arrest or delay in G1. Our genetic observations suggest that the role of p53 in activating the SCF^Ago^ for the downregulation of CycE is at play (65, 66). This p53-dependent response may function to promote the repair of nuclear damage before cells advance in the cell cycle (Fig 7F). Failure in this mechanism would increase cell death, which would be consistent with the enhanced developmental defects we observed.

We discovered that the apoptotic response to sporadic post-mitotic NR defects is crucial to tissue development. Our observation that blocking apoptosis with P35 in this context, rather than increasing wing size through the persistence of defective cells, caused a marked decrease in wing size, was striking to us. Previous studies showed that apoptotic cells in proliferating tissues produce signals (Decapentaplegic and Wingless) that induce a compensatory proliferation that promotes tissue homeostasis (75–77). Therefore, blocking apoptosis may prevent cells born with defective nuclei from triggering compensatory proliferation to promote the development of a normal-size tissue. Alternatively, the persistence of defective cells may be harmful to tissue development. Cells that enter apoptosis but fail to complete it, termed *undead* cells, are thought to overproduce proliferative signals that interfere with tissue development (75, 77). However, we think such an effect is unlikely to account for the enhancement of the small wing phenotype we observed when blocking apoptosis. This is because we found that blocking apoptosis at the level of Dronc or upstream (by expression of Diap1 or depletion of Hid) also enhanced the small wing phenotype, and it is known that the production of proliferative signals requires Dronc (78, 79). Thus, our results suggest that apoptosis upon NR defects may serve the dual purpose of eliminating injured cells while stimulating compensatory proliferation allowing their replacement.

In summary, we propose that cells born with structural nuclear defects can follow at least two different fates. In some cases, they may repair nuclear damage following a p53-dependent cell cycle arrest. In other cases, apoptosis functions as a safeguard mechanism that eliminates defective cells and promotes normal tissue development. The relative contributions of these mechanisms in different developmental contexts should be explored. Interestingly, p53 was shown to promote compensatory proliferation following apoptosis in imaginal discs (78), suggesting that both mechanisms may be connected. In addition, it would be interesting to investigate to what extent similar responses function in mammals.

## MATERIALS AND METHODS

### Plasmids, transfections and cell lines

Plasmids used in this study were generated using the Gateway recombination system (Thermo Fisher Scientific). The cDNA of each gene was cloned into the pDONOR221 vector and sequenced before being recombined into the destination vector downstream of the inducible metallothionein promoter (pMT) or the constitutive Actin 5C promoter (pAc5). The following expression plasmids were generated: pAc5-Flag-BAF, pMT-GFP-BAF, pMT-GFP-BAF^3A^, pAc5-mcherry-Tubulin, pAc5-Lamin-GFP. Point mutations were generated using QuickChange Lighting Site-Directed Mutagnenesis Kit (Agilent, #210518) according to the manufacturer’s instructions.

To generate stable cell lines, D-Mel (d.mel-2) cells were transfected using X-tremeGene HP DNA transfection reagent (#06366236001, Roche). Two days after transfection, cells were selected with 20 μg/ml blasticidin for 6 passages. While inducible pMT-based plasmids contained the blasticidin resistance gene, pAc5-based vectors were co-transfected with pCoBlast to render cells resistant to blasticidin. All the cell lines were maintained in the Express Five medium supplemented with penicillin, streptomycin, and glutamine. Cell lines expressing pMT constructs were induced with 300 μM CuS0_4_ at least 16 hours before experiments. Cells expressing GFP-BAF + mCherry-Tubulin, and cells expressing H2A-RFP + Lamin-GFP were described elsewhere (37, 80).

For knock-down experiments with RNA interference, dsRNAs were generated from PCR amplicons using a T7 RiboMAX kit (Promega). dsRNA against the bacterial kanamycine resistance gene (KAN) was used as a non-target (NT) control. 20 μg of dsRNA was transfected into 1x10^6^ D-Mel cells using Transfast transfection reagent (#E2431, Promega) according to the manufacturer’s protocol. Cells were then collected and analyzed by immunoblotting, immunofluorescence or live cell imaging.

### Fly genetics

Flies were cultured in a standard *Drosophila* agar medium at 25°C. Oregon R was used as wild-type (WT) reference and the fly lines for RNAi were provided by the Bloomington *Drosophila* Stock Center (BDSC) or Vienna Drosophila Resource Center (VDRC). All Fly strains used in this study and their sources are listed in Table S1. All crosses were also carried out at the indicated temperature (25°C, 22°C or 18°C) with 60-70% humidity. Expression of UAS transgenes in the developing wing pounch was done with the *Nubbin-GAL4* driver (*Nub-Gal4*). The selection of larvae of the desired genotypes was done by avoiding the Tb marker provided on balancer third chromosomes. To select for insertions or mutations on the second chromosomes, the *T(2;3)TSTL14, SM5: TM6B Tb^1^* pair of balancer second and third chromosomes was used. To facilitate experiments, chromosomes combining *Nub-Gal4, Ankle2 RNAi* (VDRC100665) and *Nub-Gal4, Lamin RNAi* (VDRC107419) were generated by recombination. To mark the region of interest (wing pouch) for the quantification of Dcp-1 staining and other phenotypes upon *Nub-Gal4*-driven RNAi depletions, *UAS-GFP.nls* (BDSC 4776) was used. To mark cells from the region of interest (wing pouch) for FACS analysis, *UAS-mCherry.nls* (BDSC 38425) was used.

### Immunohistochemistry in wing discs

Wandering third instar larvae were dissected in Express Five medium and fixed with 4% paraformaldehyde, 1 mM CaCl_2_ in PBS for 20 min at room temperature. Following fixation, wing discs were permeabilized and blocked in PBS, 5% BSA, 0.2% Triton for 15 min, and subsequently incubated with primary antibodies overnight at 4°C. After three washes with PBS, 0.2% Triton (PBST 0.2%), wing discs were incubated with secondary antibodies and DAPI in PBST 0.2% for with agitation 1 h at room temperature in the dark. Following three washes of 5 min each with PBST 0.2%, wing discs were isolated from the larval tissues and mounted in Moviol. Primary antibodies used were as follows: anti-cleaved Dcp-1 from rabbit (#9578, Cell signaling Technology. 1:500,), anti-Lamin from mouse (Developmental Studies Hybridoma Bank ADL84.12 deposited by P. A. Fisher, 1:500), anti-NPC from mouse (Ab24609, Abcam. 1:200), anti-Otefin from rabbit (custom made by Thermo Fisher Scientific 1:200). Anti-phospho-Histone H3 (pHH3) from rabbit (EMD Millipore, 1:200), anti-Histone pH2AvD from rabbit (#601-401-914S, Rockland, 1:200). Secondary antibodies used were purchased from Thermo Fisher Scientific and were as follows: Alexa Fluor 555 anti-rabbit (1:200), Alexa Fluor 647 anti-mouse (1:200).

### Imaginal wing disc TUNEL staining

Terminal deoxynucleotidyl transferase dUTP nick end labeling (TUNEL) assay in imaginal wing discs was performed using the ApopTag red *in situ* apoptosis detection Kit (#S7165, Sigma-Aldrich), following the manufacturer’s procedure. Briefly, dissected larval wing discs were fixed in 4% paraformaldehyde in PBS for 20 min at room temperature, then washed twice in PBST 0.2% for 5 min. A post-fixation was done in precooled ethanol: acetic acid 2:1 for 5 min at -20°C. After three washes with PBST 0.2%, larval tissues were incubated in 75 μl of Equilibration buffer (90416) for 10 min at room temperature. After completely removing the Equilibration buffer, 55 μl of freshly prepared Working strength TdT reaction buffer was added into larval tissues and incubated for 1 hour at 37°C in a humidified chamber. Shortly after applying stop/wash buffer (90419) and 3 washes with PBST 0.2%, tissues were incubated in the dark with anti-Digoxigenin conjugate working buffer (53% v/v Blocking solution (90425) and 47% v/v anti-digoxigenin conjugate (90429)) with DAPI for 30 min at room temperature and mounted in moviol. For TdT end- and Dcp-1 double-labeling, the anti-Dcp-1 antibody was prepared with the blocking solution (90425). Blocking and primary antibody staining were carried out as described for the immunohistochemistry after applying stop/wash buffer. The secondary antibody was diluted with the anti-Digoxigenin conjugate working buffer.

### Imaginal wing disc EdU labeling

Imaginal wing discs from third instar larvae were dissected in Express Five medium at room temperature and incubated for 1 hour with 10 μM EdU. Wing discs were then fixed and stained, according to standard protocol (Click-iT EdU cell proliferation Kit for imaging, Alexa Fluor 488, Thermo Fisher Scientific, #C10337). After removing the cocktail reaction buffer, three short washes were done and additional stanning was applied.

### Western blotting and immunofluorescence in cells in culture

For Western blots, cells were resuspended in PBS containing protease inhibitors and 1 volume of 2X Laemmli buffer (S3401-10VL, Sigma) was added before heating at 95°C for 2 min. Extracts were then electrophoresed on 12% Tris-acrylamide SDS-PAGE gel and blotted onto a PVDF membrane (#1620177, Bio-Rad). Membranes were blocked with 5% milk solution for 1 hour at room temperature and probed with primary antibody overnight at 4°C. Peroxidase-conjugated anti-rabbit secondary antibodies from goat (1:5000, 111-035-008, Jackson ImmunoResearch) or anti-mouse from goat (1:5000, 115-035-003, Jackson ImmunoResearch) were used for primary antibody detection. To visualize protein phosphorylation levels, Phos-tag (#300-93523, FUJIFILM Wako Chemicals) was added into 8% Tris-acrylamide SDS-PAGE gel. After electrophoresis, the gel was soaked in transfer buffer with 1 mM EDTA for 10 min and then in transfer buffer without EDTA for another 10 min before transferring onto a PDVF membrane. Primary antibodies used for WB were as follows: anti-α-Tubulin DM1A from mouse (#T6199, Sigma, 1:5000), anti-Lamin Dm0 from mouse (Developmental Studies Hybridoma Bank ADL84.12 deposited by P. A. Fisher, 1:1000), anti-BAF (custom made by Thermo scientific, 1:1000), anti-Ankle2 (custom made by Thermo Fisher Scientific 1:1000), anti-GFP from rabbit (TP401, Torrey Pines, 1:5000). The blots were visualized using Clarity Western ECL substrate (170-5060, Bio-Rad) by the ChemiDoc XRS+ system (Bio-Rad).

For immunofluorescence, D-Mel Cells were washed 2 times with PBS before fixation on coverslips with 4% formaldehyde for 20 min at room temperature. After fixation, cells were permeabilized and blocked in PBS containing 0.1% triton X-100 and 1% BSA (PBSBT) for 15 min. Cells were then incubated with primary antibodies diluted in PBSBT for 2 hours at room temperature, washed three times in PBS containing 0.1% triton X-100 (PBST) and incubated with secondary antibodies and DAPI diluted in PBSBT for 1 hour at room temperature. Coverslip were washed 3 times in PBST before being mounted in Vectashield medium (Vector). Primary antibodies used were as follows: anti-Flag from mouse (#F1804, Sigma-Aldrich, 1:2000), anti-Lamin Dm0 from mouse (Developmental Studies Hybridoma Bank ADL84.12 deposited by P. A. Fisher, 1:500), Alexa Fluor 488 phalloidin (#A12379, Invitrogen, 1:1000), anti-Dcp-1 from rabbit (#9578, Cell signaling Technology, 1:200). Secondary antibodies used were purchased from Thermo Fisher Scientific and were as follows: Alexa Fluor 555 anti-rabbit (1:200), Alexa Fluor 488 anti-rabbit (1:200), Alexa Fluor 488 anti-mouse (1:200), Alexa Fluor 647 anti-mouse (1:200).

### Microscopy and quantification

Live imaging in D-Mel cells was performed using a Spinning-disk confocal system (Yokogawa CSU-X1 5000) mounted on a fluorescence microscope (Zeiss Axio Observer Z1). Cells were plated into a LabTek II chambered coverglass (#155409, Thermo Fisher Scientific) for at least 2 hours before filming. To monitor GFP-BAF and Lamin-GFP levels over time, GFP signals for each time point at the reassembling nuclei (circular areas) was quantified directly with the Zen software.

All Images of imaginal wing discs were taken using the scanning confocal microscope Zeiss Leica SP8. for each imaginal wing disc, 15focal planes were taken at 0.35 μM intervals and analyzed using Fiji software. Six to 8 wing discs in each condition were used for the quantification of Dcp-1 staining. Briefly, a Z-sum intensity projection was generated for each image and the GFP positive area was defined as the region of interest. Three regions out of the zone of interest in the disc were randomly chosen and the mean intensity of these zones (background) was calculated. the specific intensity of Dcp-1 in the region of interest was obtained by subtraction of the background.

The scanning confocal microscope Zeiss LSM880 was used to take images of fixed D-Mel cells and cells from fixed imaginal wing discs (higher magnification than entire discs). Multiple focal planes at 0.16 μm intervals were obtained and subsequently treated with AiryScan software. For quantifications of fixed cells, mean intensity of fluorescence was measured directly with ZEN software in a single in-focus plane to monitor Flag-BAF fluorescence in the nucleus, the cytoplasm or at the nuclear periphery. To quantify the number of cells with nuclear phenotypes of interest, maximum-intensity projections were interest were labeled by hand and the cell counter tool in Fiji software was used to obtain cell numbers.

All images of adult wings were taken using a stereo microscope. Fifteen wings from adult females were analyzed for each condition using Fiji software and the wing size was quantified with the area measurement tool.

### Fluorescence Activated Cell Sorting

At least 30 wing discs form third instar larvae were dissected in Express 5 medium at room temperature and washed twice for 3 min in PBS. Cells were dissociated by incubating wing discs in 10x trypsin-EDTA solution (#T4174, Sigma-Aldrich,) in the presence of the DNA dye Hoechst 33342 (#10337G, Invitrogen, 0,5ug/ml) for 3 hours with gentle agitation. Cells were filtered and analyzed with a Yeti Cell Analyzer and Flowjo software.

## ACKNOWLEDGEMENTS

This work was funded by a Project Grant from the Canadian Institutes of Health Research (CIHR) to VA (175132). JL and HM received studentships from the Fonds de Recherche du Québec – Santé. We thank Christian Charbonneau and Annie Gosselin for precious help with the microscopy and FACS, respectively. We thank Caroline Baril and David Hipfner for sharing fly lines, for technical help and for helpful discussions. We thank Pamela Geyer and Talila Volk for sharing fly lines.

## AUTHOR CONTRIBUTIONS

JL, LJ, HM and XW conducted the experiments. JL and VA designed the experiments and wrote the paper.

**Video 1. Complement to Figure 1E. Mitosis in a cell expressing GFP-BAF (green) and mCherry-Tubulin (magenta) after Non-Target RNAi.**

**Video 2. Complement to Figure 1E. Mitosis in a cell expressing GFP-BAF (green) and mCherry-Tubulin (magenta) after Ankle2 RNAi.**

**Video 3. Complement to Figure 2E. Mitosis in a cell expressing Lamin-GFP (green) and H2A-RFP (magenta) after Non-Target RNAi.**

**Video 4. Complement to Figure 2E. Mitosis in a cell expressing Lamin-GFP (green) and H2A-RFP (magenta) after Ankle2 RNAi.**

**Table S1. *Drosophila* strains used in this study.** The transgenes, mutant alleles, sources and complete genotypes are indicated.

---

**Figure S1.**
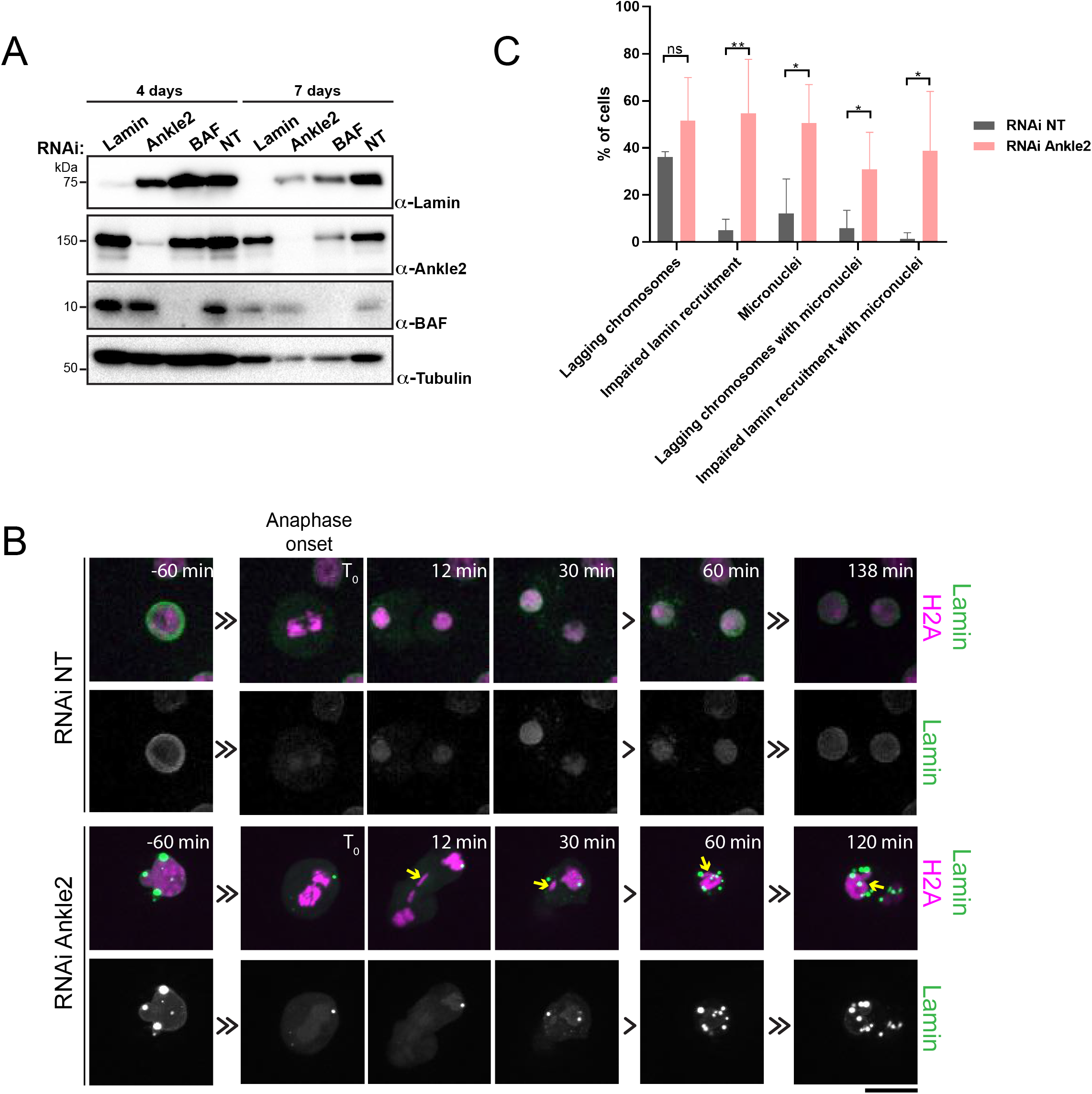
Complement to Figure 2 - Ankle2, BAF and Lamin are required for nuclear reassembly in cells in culture. **A.** Western blots showing RNAi depletion of Ankle2, BAF and Lamin 4- or 7-days post transfection with the indicated dsRNA. NT: Non-Target. **B.** Depletion of Ankle2 causes post-mitotic micronucleation. D-Mel cells expressing H2A-RFP and GFP-Lamin were transfected with the indicated dsRNA for 4 days and mitoses were filmed. Arrows: a lagging chromosome in telophase becomes a micronucleus. Scale bar: 5 μm. **C.** Quantification of nuclear defects observed by live imaging after Ankle2 RNAi (or NT control) as in B. *p < 0.05, **p < 0.01, ns: non-significant from unpaired t-tests.

**Figure S2.**
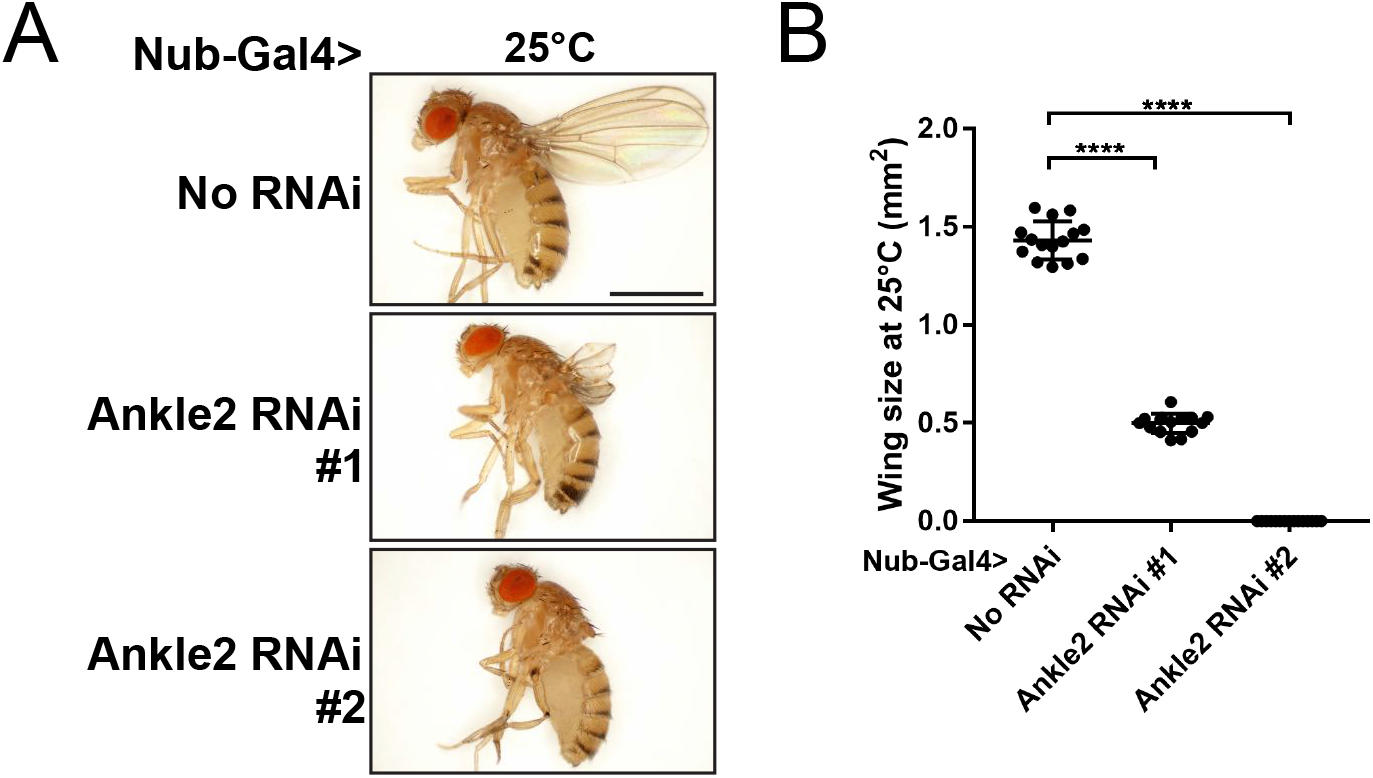
Wing development defects caused by the depletion of Ankle2 using two RNAi. Ankle2 RNAi was driven by Nub-Gal4 at 25°C using lines VDRC100655 (Ankle2 RNAi line #1) and BDSC77437 (Ankle2 RNAi line #2). **A.** Example images of adult flies. Scale bar: 1 mm. **B.** Quantification of wing sizes at 25°C. ****p < 0.0001 from unpaired t-tests with Welch’s correction.

**Figure S3.**
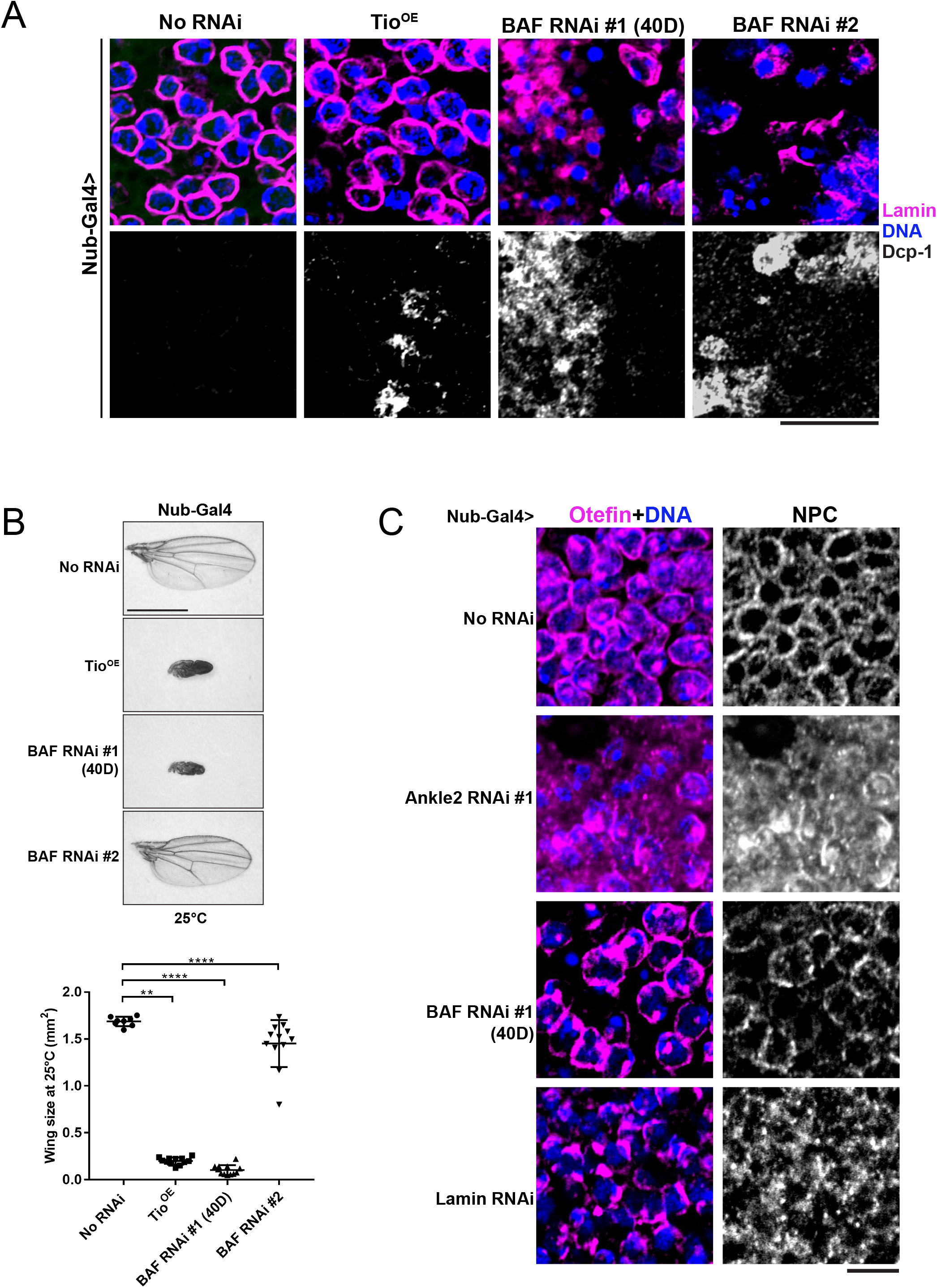
Complement to Figure 3 – RNAi Depletion of Ankle2, BAF or Lamin during wing development causes nuclear defects and adult wing defects. **A.** Nuclear defects in wing discs resulting from driving BAF RNAi from two lines (VDRC 103013, left and BDSC36108, right) with Nub-Gal4 at 25°C. Both BAF RNAi constructions induce Lamin mislocalization and DNA foci devoid of Lamin. The VDRC103013 line has the BAF RNAi construction inserted at cytolocation 40D which was shown to also lead to overexpression of Tio. A control line with a UAS element alone inserted at the same site (VDRC60101, Tio^OE^) does not result in similar nuclear defects. Therefore, these defects are specific to BAF depletion for both BAF RNAi lines. However, the Tio^OE^ line results in some level of apoptosis revealed by Dcp-1 staining. Scale bar: 10 μm. **B.** Adult wing defects resulting from driving the two BAF RNAi insertions with Nub-Gal4 at 25°C. Top: Examples of adult wings of the indicated genotypes. Line VDRC103013 results in a more pronounced phenotype but the control Tio^OE^ line results in a similar small wing phenotype. Therefore, the adult wing phenotype with line VDRC103013 is not specific to BAF depletion. Scale bar: 1 mm. Bottom: Quantification of wing sizes at 25°C. **p < 0.01, ****p < 0.0001, ns: non-significant from unpaired t-tests with Welch’s correction. **C.** RNAi Depletion of Ankle2 (VDRC100655), BAF (VDRC103013) or Lamin results in mislocalization of Otefin and NPC proteins. All constructions were driven with Nub-Gal4 at 25°C. Scale bar: 5 μm.

**Figure S4.**
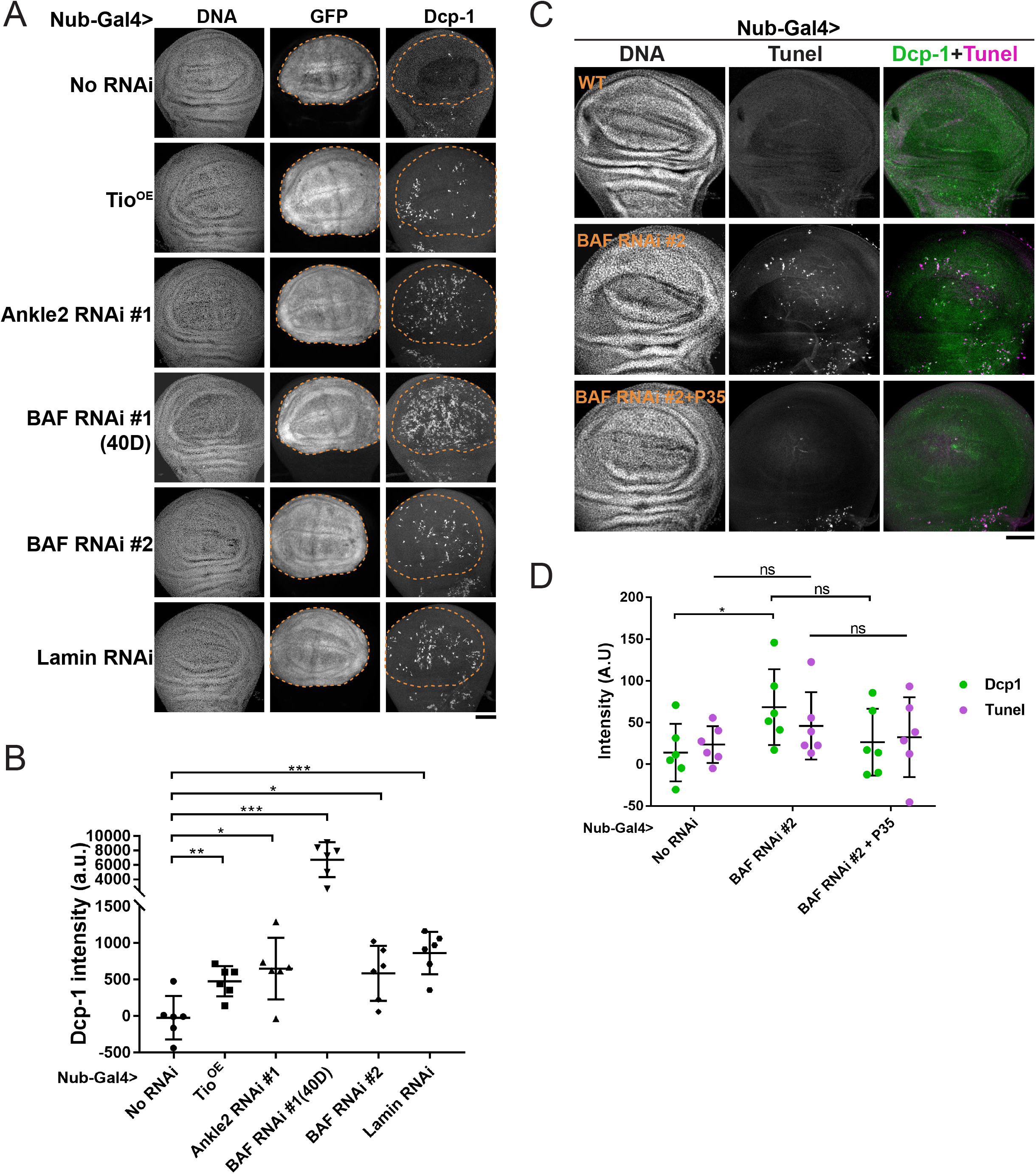
Complement to Figure 4 – RNAi depletion of Ankle2, BAF or Lamin causes apoptosis. **A.** The indicated RNAi constructions were induced in the wing pouch by Nub-Gal4 at 25°C. In parallel, UAS-GFP was used as a marker of the region of interest (wing pouch, dotted lines). Wing discs were analyzed by immunofluorescence against cleaved Dcp-1 and stained for DNA (DAPI). **B.** Quantification of the Dcp-1 intensities measured in the wing pouch region of individual wing discs of the indicated genotypes as in A. **C.** Detection of apoptosis by TUNEL (magenta) and simultaneous immunofluorescence for cleaved Dcp-1 (green) in wing discs depleted of BAF. Expression of P35 abrogates apoptosis. **D.** Quantifications of TUNEL and Dcp-1 signals in wing discs of the indicated genotypes. In all panels: BAF RNAi #1 (40D): VDRC103013; BAF RNAi #2: BDSC36108; Ankle2 RNAi #1: VDRC100655. All Scale bars: 50 μm. All error bars: S.D. *p<0.05, **p < 0.01, *** p < 0.001 ns: non-significant from unpaired t-tests with Welch’s correction.

**Figure S5.**
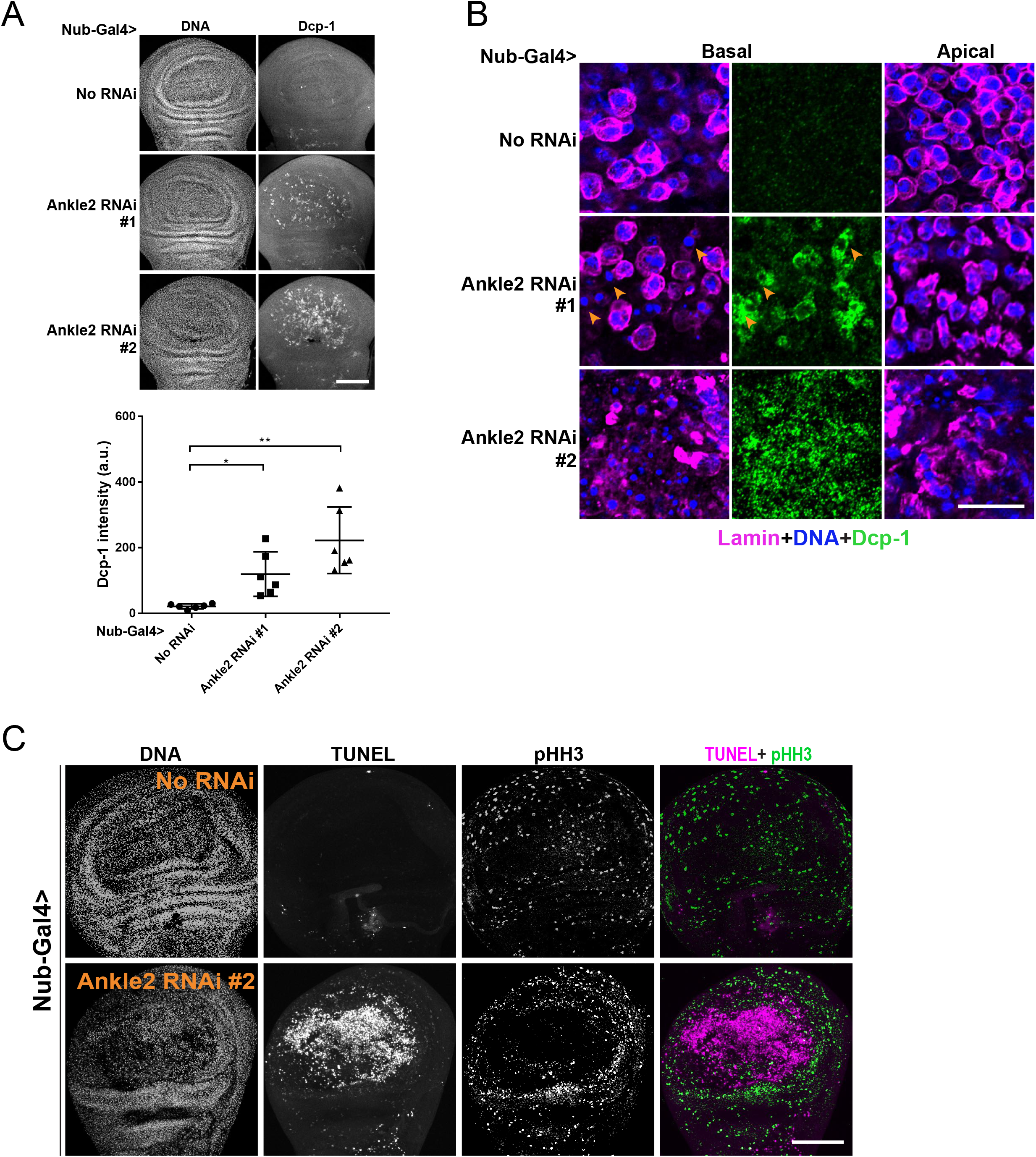
Complement to Figure 4 – RNAi depletion of Ankle2 using two RNAi constructions causes apoptosis in wing discs. **A.** Ankle2 RNAi was driven by Nub-Gal4 at 25°C using lines VDRC100655 (Ankle2 RNAi line #1) and BDSC77437 (Ankle2 RNAi line #2). Top: Cleaved Dcp-1 staining in wing discs of the indicated genotypes. Bottom: Quantification of Dcp-1 signals. All error bars: S.D. *p<0.05, **p < 0.01, from unpaired t-tests with Welch’s correction. **B.** Nuclear defects and apoptosis are revealed by immunofluorescence against Lamin, DAPI and cleaved Dcp-1. Arrowheads: hypercondensed DNA in apoptotic cells. Scale bar: 10 μm. **C.** phospho-Histone H3 (pHH3) staining suggests compensatory cell proliferation around the area of the wing disc where Ankle2 is depleted (line BDSC77437) and apoptosis occurs (detected by TUNEL). All scale bars for imaginal wing discs: 50 μm.

**Figure S6.**
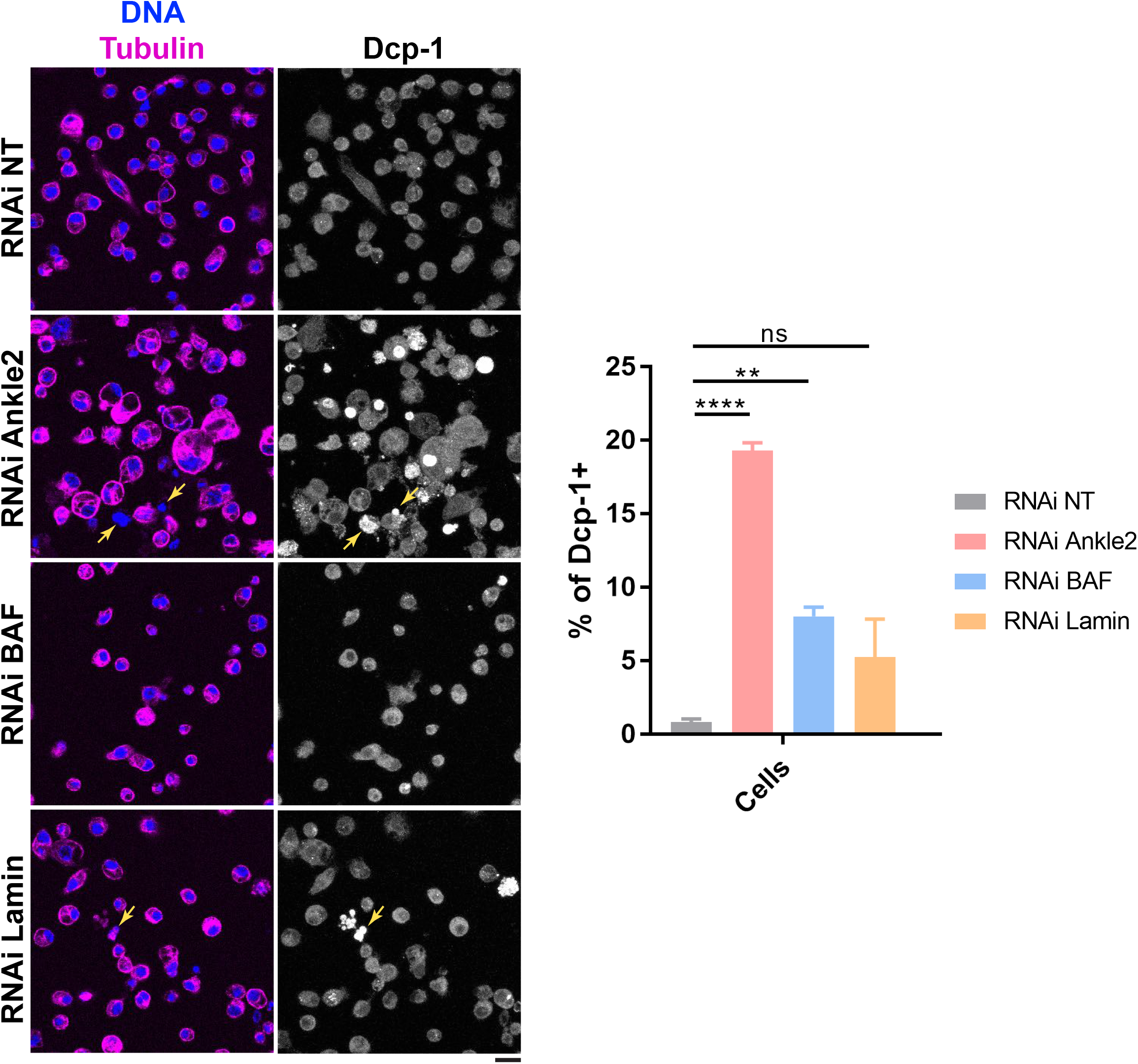
RNAi depletion or Ankle2 or BAF results in apoptosis in D-Mel cells in culture. D-Mel cells were transfected with the indicated dsRNA and analyzed by immunofluorescence after 4 days. Left: Examples of images. Right: Quantification of Dcp-1 positive (Dcp-1+) cells of indicated conditions. Arrows: apoptotic cells with hypercondensed DNA. Scale bar: 20 μm. All error bars: S.D. **p < 0.01, **** p < 0.0001, ns: non-significant from unpaired t-tests.

**Figure S7.**
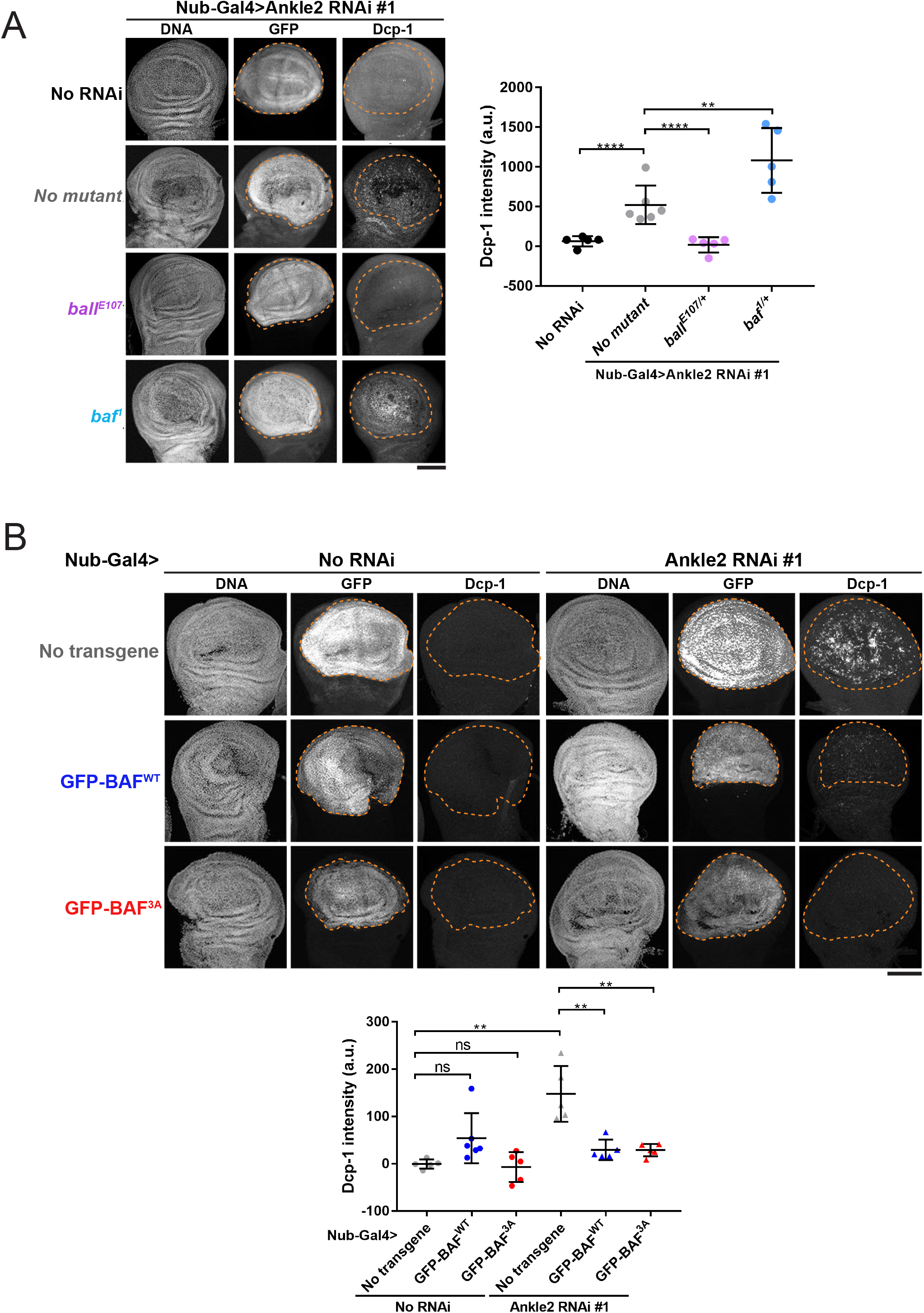
Disruption of Ankle2 function in BAF regulation during wing development causes apoptosis. **A.** Mutation in *baf* enhances, while mutation in *ball* suppresses, apoptosis resulting from Ankle2 depletion. Left: Examples of wing discs of the indicated genotypes after induction of Ankle2 RNAi using Nub-Gal4 at 25°C. In parallel, UAS-GFP was used as a marker of the region of interest (wing pouch, inside dotted line). Right: Quantification of Dcp-1 signals in the wing pouch. **B.** Overexpression of GFP-BAF^3A^ or GFP-BAF^WT^ rescues apoptosis resulting from Ankle2 depletion. Left: Examples of wing discs of the indicated genotypes at 25°C. Right: Quantification of Dcp-1 signals. Analysis was done as in A. In all experiments, Ankle2 depletion was done using line VDRC100655. All scale bars: 50 μm. All error bars: S.D. **p < 0.01, ***p < 0.001, from unpaired t-tests with Welch’s correction.

**Figure S8.**
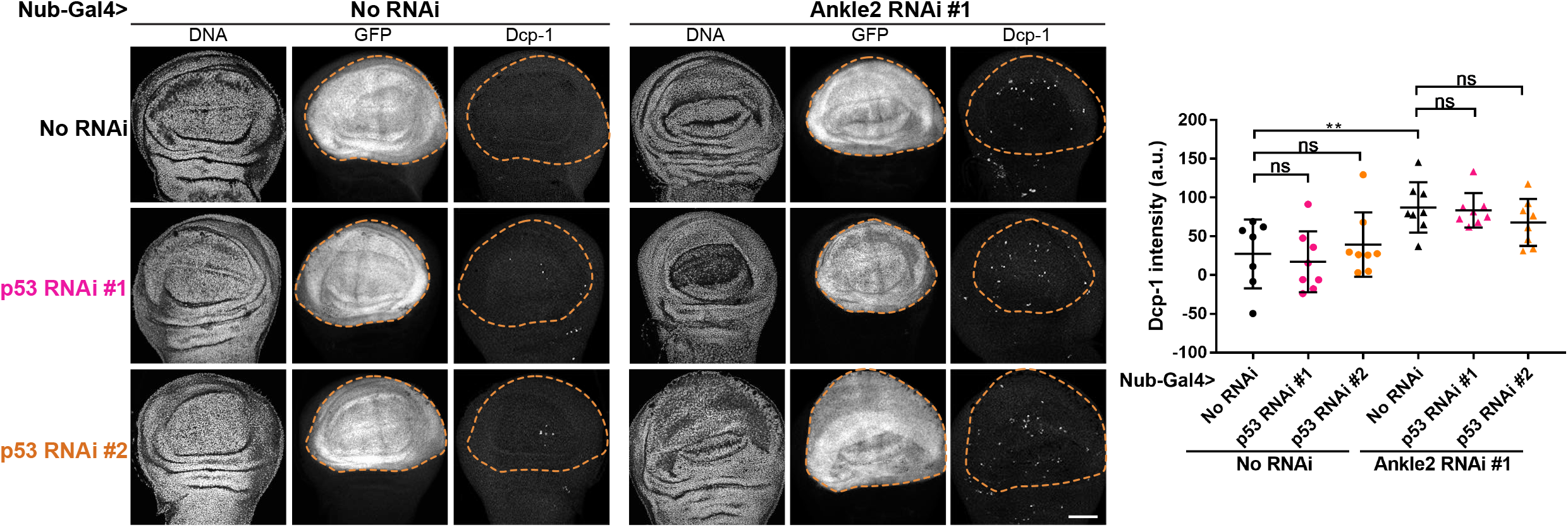
Complement to Figures 6 - p53 is not required for apoptosis in response to nuclear reassembly defects. **A.** Depletion of p53 does not block apoptosis resulting from Ankle2 depletion (line VDRC100655). RNAi depletions of p53 were achieved with lines VDRC38235 (p53 RNAi #1) and VDRC10692 (p53 RNAi #2). Left: Examples of wing discs of the indicated genotypes after induction of Ankle2 RNAi using Nub-Gal4 at 25°C, UAS-GFP is used as a maker for the region of interest (pouch area). Right: Quantifications of Dcp-1 signals in the wing pouch. Scale bar: 50 μm. All error bars: S.D. **p < 0.01, ns: non-significant from unpaired t-tests with Welch’s correction.

**Figure S9.**
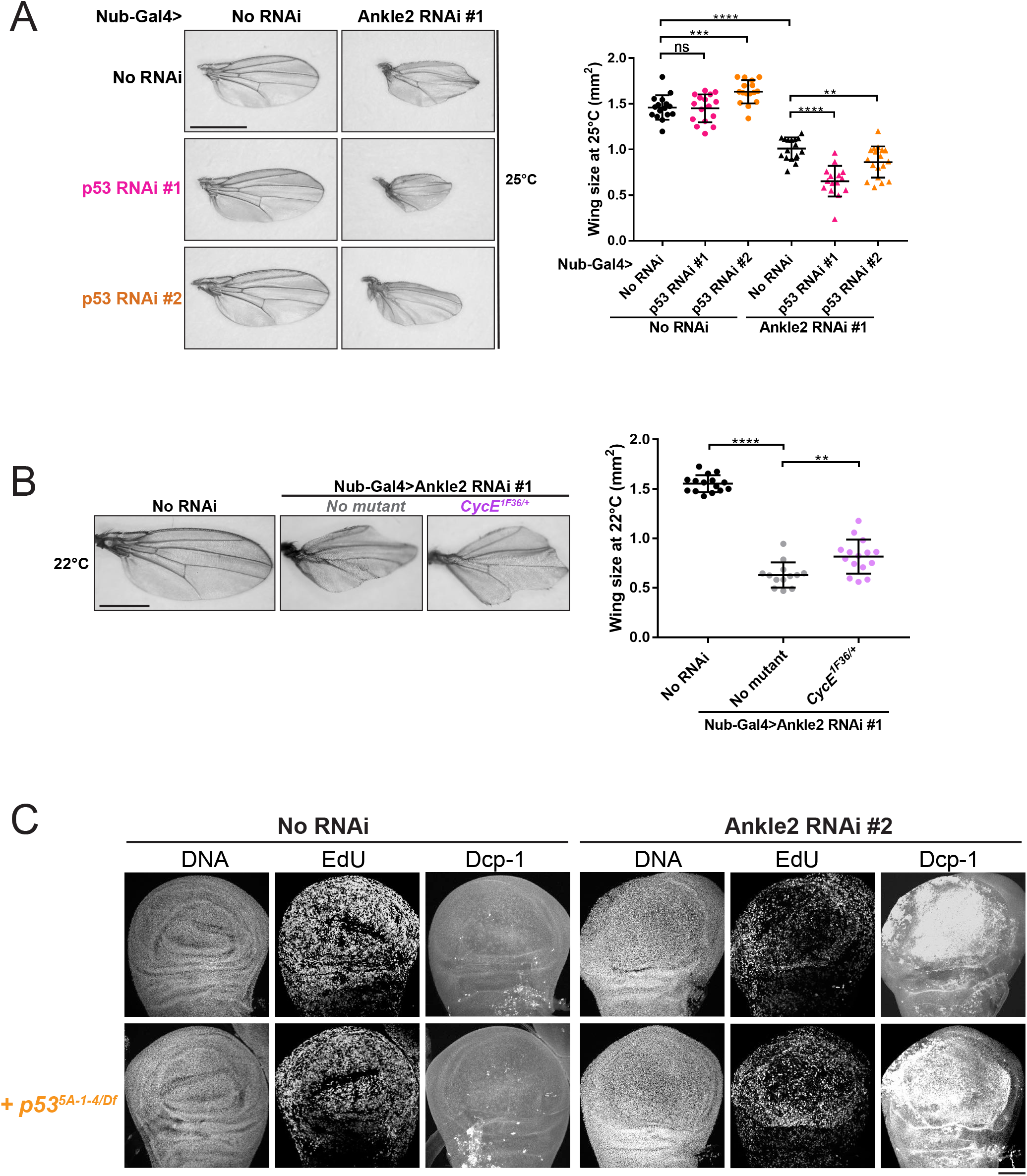
Complement to Figure 7 - A p53-dependent response promotes tissue development when nuclear reassembly is compromised. **A.** Depletions of p53 enhance the small wing phenotype resulting from Ankle2 depletion. Left: Examples of adult wing of the indicated genotypes after inducing RNAi using Nub-Gal4 at 25°C. Right: Quantifications of wing sizes at 25°C. Scale bar: 1 mm. **B.** Mutation in *cycE* enhance the small wing phenotype resulting from Ankle2 depletion. Left: Examples of adult wing of the indicated genotypes at 22°C and 18°C. Right: Quantifications of wing sizes. Scale bar: 1 mm. In panels A-B, Ankle2 RNAi line VDRC100655 was used. All error bars: S.D. **p < 0.01, *** p < 0.001, **** p < 0.0001 ns: non-significant from unpaired t-tests with Welch’s correction. **C.** Decreased population of EdU+ cells resulting from Ankle2 depletion (line BDSC77437) in wing discs is rescued by inactivation of *p53*. Wings discs of indicated genotypes were incubated with EdU for 1 hour and then co-stained with DAPI (DNA) and cleaved Dcp-1. Scale bar: 50 μm.

## REFERENCES

1. Morgan DO. The Cell Cycle: Principles of Control. London: New Science Press; 2007. 297 p.

2. Alberts B, Johnson A, Lewis J, Morgan D, Raff M, Roberts K, et al. Molecular Biology of the Cell. Edition S, editor. United States: W. W. Norton & Co.; 2015. 1464 p.

3. Schellhaus AK, De Magistris P, Antonin W. Nuclear Reformation at the End of Mitosis. J Mol Biol. 2016;428(10 Pt A):1962–85.

4. Liu S, Pellman D. The coordination of nuclear envelope assembly and chromosome segregation in metazoans. Nucleus. 2020;11(1):35–52.

5. Hintzsche H, Hemmann U, Poth A, Utesch D, Lott J, Stopper H, et al. Fate of micronuclei and micronucleated cells. Mutat Res. 2017;771:85–98.

6. Kamikawa Y, Imaizumi K. Advances in understanding the mechanisms of repairing damaged nuclear envelop. J Biochem. 2022;171(6):609–17.

7. de Leeuw R, Gruenbaum Y, Medalia O. Nuclear Lamins: Thin Filaments with Major Functions. Trends Cell Biol. 2018;28(1):34–45.

8. Nazer E. To be or not be (in the LAD): emerging roles of lamin proteins in transcriptional regulation. Biochem Soc Trans. 2022;50(2):1035–44.

9. Liu SY, Ikegami K. Nuclear lamin phosphorylation: an emerging role in gene regulation and pathogenesis of laminopathies. Nucleus. 2020;11(1):299–314.

10. Donnaloja F, Carnevali F, Jacchetti E, Raimondi MT. Lamin A/C Mechanotransduction in Laminopathies. Cells. 2020;9(5).

11. Maciejowski J, Hatch EM. Nuclear Membrane Rupture and Its Consequences. Annu Rev Cell Dev Biol. 2020;36:85–114.

12. Lukasova E, Kovarik A, Kozubek S. Consequences of Lamin B1 and Lamin B Receptor Downregulation in Senescence. Cells. 2018;7(2).

13. Dubik N, Mai S. Lamin A/C: Function in Normal and Tumor Cells. Cancers (Basel). 2020;12(12).

14. Irianto J, Pfeifer CR, Ivanovska IL, Swift J, Discher DE. Nuclear lamins in cancer. Cell Mol Bioeng. 2016;9(2):258–67.

15. Hatch EM, Fischer AH, Deerinck TJ, Hetzer MW. Catastrophic nuclear envelope collapse in cancer cell micronuclei. Cell. 2013;154(1):47–60.

16. Guo X, Dai X, Wu X, Cao N, Wang X. Small but strong: Mutational and functional landscapes of micronuclei in cancer genomes. Int J Cancer. 2021;148(4):812–24.

17. Zhang CZ, Spektor A, Cornils H, Francis JM, Jackson EK, Liu S, et al. Chromothripsis from DNA damage in micronuclei. Nature. 2015;522(7555):179-84.

18. Hopfner KP, Hornung V. Molecular mechanisms and cellular functions of cGAS-STING signalling. Nat Rev Mol Cell Biol. 2020;21(9):501–21.

19. Decordier I, Dillen L, Cundari E, Kirsch-Volders M. Elimination of micronucleated cells by apoptosis after treatment with inhibitors of microtubules. Mutagenesis. 2002;17(4):337–44.

20. Guo X, Ni J, Liang Z, Xue J, Fenech MF, Wang X. The molecular origins and pathophysiological consequences of micronuclei: New insights into an age-old problem. Mutat Res. 2019;779:1–35.

21. Sears RM, Roux KJ. Diverse cellular functions of barrier-to-autointegration factor and its roles in disease. J Cell Sci. 2020;133(16).

22. Samwer M, Schneider MWG, Hoefler R, Schmalhorst PS, Jude JG, Zuber J, et al. DNA Cross-Bridging Shapes a Single Nucleus from a Set of Mitotic Chromosomes. Cell. 2017;170(5):956–72 e23.

23. Zheng R, Ghirlando R, Lee MS, Mizuuchi K, Krause M, Craigie R. Barrier-to-autointegration factor (BAF) bridges DNA in a discrete, higher-order nucleoprotein complex. Proc Natl Acad Sci U S A. 2000;97(16):8997–9002.

24. Umland TC, Wei SQ, Craigie R, Davies DR. Structural basis of DNA bridging by barrier-to-autointegration factor. Biochemistry. 2000;39(31):9130–8.

25. Lee MS, Craigie R. A previously unidentified host protein protects retroviral DNA from autointegration. Proc Natl Acad Sci U S A. 1998;95(4):1528–33.

26. Haraguchi T, Koujin T, Segura-Totten M, Lee KK, Matsuoka Y, Yoneda Y, et al. BAF is required for emerin assembly into the reforming nuclear envelope. J Cell Sci. 2001;114(Pt 24):4575–85.

27. Furukawa K. LAP2 binding protein 1 (L2BP1/BAF) is a candidate mediator of LAP2-chromatin interaction. J Cell Sci. 1999;112 (Pt 15):2485–92.

28. Lee KK, Haraguchi T, Lee RS, Koujin T, Hiraoka Y, Wilson KL. Distinct functional domains in emerin bind lamin A and DNA-bridging protein BAF. J Cell Sci. 2001;114(Pt 24):4567–73.

29. Mansharamani M, Wilson KL. Direct binding of nuclear membrane protein MAN1 to emerin in vitro and two modes of binding to barrier-to-autointegration factor. J Biol Chem. 2005;280(14):13863–70.

30. Barton LJ, Soshnev AA, Geyer PK. Networking in the nucleus: a spotlight on LEM-domain proteins. Curr Opin Cell Biol. 2015;34:1–8.

31. Nichols RJ, Wiebe MS, Traktman P. The vaccinia-related kinases phosphorylate the N’ terminus of BAF, regulating its interaction with DNA and its retention in the nucleus. Mol Biol Cell. 2006;17(5):2451–64.

32. Birendra K, May DG, Benson BV, Kim DI, Shivega WG, Ali MH, et al. VRK2A is an A-type lamin-dependent nuclear envelope kinase that phosphorylates BAF. Mol Biol Cell. 2017;28(17):2241–50.

33. Asencio C, Davidson IF, Santarella-Mellwig R, Ly-Hartig TB, Mall M, Wallenfang MR, et al. Coordination of kinase and phosphatase activities by Lem4 enables nuclear envelope reassembly during mitosis. Cell. 2012;150(1):122–35.

34. Snyers L, Erhart R, Laffer S, Pusch O, Weipoltshammer K, Schofer C. LEM4/ANKLE-2 deficiency impairs post-mitotic re-localization of BAF, LAP2alpha and LaminA to the nucleus, causes nuclear envelope instability in telophase and leads to hyperploidy in HeLa cells. Eur J Cell Biol. 2018;97(1):63–74.

35. Apridita Sebastian W, Shiraishi H, Shimizu N, Umeda R, Lai S, Ikeuchi M, et al. Ankle2 deficiency-associated microcephaly and spermatogenesis defects in zebrafish are alleviated by heterozygous deletion of vrk1. Biochem Biophys Res Commun. 2022;624:95–101.

36. Furukawa K, Sugiyama S, Osouda S, Goto H, Inagaki M, Horigome T, et al. Barrier-to-autointegration factor plays crucial roles in cell cycle progression and nuclear organization in Drosophila. J Cell Sci. 2003;116(Pt 18):3811–23.

37. Mehsen H, Boudreau V, Garrido D, Bourouh M, Larouche M, Maddox PS, et al. PP2A-B55 promotes nuclear envelope reformation after mitosis in Drosophila. J Cell Biol. 2018;217(12):4106–23.

38. Lancaster OM, Cullen CF, Ohkura H. NHK-1 phosphorylates BAF to allow karyosome formation in the Drosophila oocyte nucleus. J Cell Biol. 2007;179(5):817–24.

39. Link N, Chung H, Jolly A, Withers M, Tepe B, Arenkiel BR, et al. Mutations in ANKLE2, a ZIKA Virus Target, Disrupt an Asymmetric Cell Division Pathway in Drosophila Neuroblasts to Cause Microcephaly. Dev Cell. 2019;51(6):713–29 e6.

40. Shah PS, Link N, Jang GM, Sharp PP, Zhu T, Swaney DL, et al. Comparative Flavivirus-Host Protein Interaction Mapping Reveals Mechanisms of Dengue and Zika Virus Pathogenesis. Cell. 2018;175(7):1931–45 e18.

41. Thomas AX, Link N, Robak LA, Demmler-Harrison G, Pao EC, Squire AE, et al. ANKLE2-related microcephaly: A variable microcephaly syndrome resembling Zika infection. Ann Clin Transl Neurol. 2022;9(8):1276–88.

42. Haraguchi T, Kojidani T, Koujin T, Shimi T, Osakada H, Mori C, et al. Live cell imaging and electron microscopy reveal dynamic processes of BAF-directed nuclear envelope assembly. J Cell Sci. 2008;121(Pt 15):2540–54.

43. Zirin JD, Mann RS. Nubbin and Teashirt mark barriers to clonal growth along the proximal-distal axis of the Drosophila wing. Dev Biol. 2007;304(2):745–58.

44. Vissers JH, Manning SA, Kulkarni A, Harvey KF. A Drosophila RNAi library modulates Hippo pathway-dependent tissue growth. Nat Commun. 2016;7:10368.

45. Fuchs Y, Steller H. Live to die another way: modes of programmed cell death and the signals emanating from dying cells. Nat Rev Mol Cell Biol. 2015;16(6):329–44.

46. Song Z, McCall K, Steller H. DCP-1, a Drosophila cell death protease essential for development. Science. 1997;275(5299):536–40.

47. Hay BA, Wolff T, Rubin GM. Expression of baculovirus P35 prevents cell death in Drosophila. Development. 1994;120(8):2121–9.

48. Callus BA, Vaux DL. Caspase inhibitors: viral, cellular and chemical. Cell Death Differ. 2007;14(1):73–8.

49. Meier P, Silke J, Leevers SJ, Evan GI. The Drosophila caspase DRONC is regulated by DIAP1. EMBO J. 2000;19(4):598–611.

50. Hawkins CJ, Yoo SJ, Peterson EP, Wang SL, Vernooy SY, Hay BA. The Drosophila caspase DRONC cleaves following glutamate or aspartate and is regulated by DIAP1, HID, and GRIM. J Biol Chem. 2000;275(35):27084–93.

51. Perez-Garijo A, Fuchs Y, Steller H. Apoptotic cells can induce non-autonomous apoptosis through the TNF pathway. Elife. 2013;2:e01004.

52. Klemm J, Stinchfield MJ, Harris RE. Necrosis-induced apoptosis promotes regeneration in Drosophila wing imaginal discs. Genetics. 2021;219(3).

53. Dos Santos A, Toseland CP. Regulation of Nuclear Mechanics and the Impact on DNA Damage. Int J Mol Sci. 2021;22(6).

54. Nagata S, Nagase H, Kawane K, Mukae N, Fukuyama H. Degradation of chromosomal DNA during apoptosis. Cell Death Differ. 2003;10(1):108–16.

55. Barton LJ, Duan T, Ke W, Luttinger A, Lovander KE, Soshnev AA, et al. Nuclear lamina dysfunction triggers a germline stem cell checkpoint. Nat Commun. 2018;9(1):3960.

56. Duan T, Kitzman SC, Geyer PK. Survival of Drosophila germline stem cells requires the chromatin-binding protein Barrier-to-autointegration factor. Development. 2020;147(9).

57. Kurzhals RL, Titen SW, Xie HB, Golic KG. Chk2 and p53 are haploinsufficient with dependent and independent functions to eliminate cells after telomere loss. PLoS Genet. 2011;7(6):e1002103.

58. Zhou L. P53 and Apoptosis in the Drosophila Model. Adv Exp Med Biol. 2019;1167:105–12.

59. Ingaramo MC, Sanchez JA, Dekanty A. Regulation and function of p53: A perspective from Drosophila studies. Mech Dev. 2018;154:82–90.

60. Smith HL, Southgate H, Tweddle DA, Curtin NJ. DNA damage checkpoint kinases in cancer. Expert Rev Mol Med. 2020;22:e2.

61. Brodsky MH, Weinert BT, Tsang G, Rong YS, McGinnis NM, Golic KG, et al. Drosophila melanogaster MNK/Chk2 and p53 regulate multiple DNA repair and apoptotic pathways following DNA damage. Mol Cell Biol. 2004;24(3):1219–31.

62. Peters M, DeLuca C, Hirao A, Stambolic V, Potter J, Zhou L, et al. Chk2 regulates irradiation-induced, p53-mediated apoptosis in Drosophila. Proc Natl Acad Sci U S A. 2002;99(17):11305–10.

63. Engeland K. Cell cycle regulation: p53-p21-RB signaling. Cell Death Differ. 2022;29(5):946–60.

64. Chen J. The Cell-Cycle Arrest and Apoptotic Functions of p53 in Tumor Initiation and Progression. Cold Spring Harb Perspect Med. 2016;6(3):a026104.

65. Mandal S, Freije WA, Guptan P, Banerjee U. Metabolic control of G1-S transition: cyclin E degradation by p53-induced activation of the ubiquitin-proteasome system. J Cell Biol. 2010;188(4):473–9.

66. Ouyang Y, Song Y, Lu B. dp53 Restrains ectopic neural stem cell formation in the Drosophila brain in a non-apoptotic mechanism involving Archipelago and cyclin E. PLoS One. 2011;6(11):e28098.

67. Samson C, Petitalot A, Celli F, Herrada I, Ropars V, Le Du MH, et al. Structural analysis of the ternary complex between lamin A/C, BAF and emerin identifies an interface disrupted in autosomal recessive progeroid diseases. Nucleic Acids Res. 2018;46(19):10460–73.

68. Holaska JM, Lee KK, Kowalski AK, Wilson KL. Transcriptional repressor germ cell-less (GCL) and barrier to autointegration factor (BAF) compete for binding to emerin in vitro. J Biol Chem. 2003;278(9):6969–75.

69. Sears RM, Roux KJ. Mechanisms of A-Type Lamin Targeting to Nuclear Ruptures Are Disrupted in LMNA- and BANF1-Associated Progerias. Cells. 2022;11(5).

70. Janssen A, Marcelot A, Breusegem S, Legrand P, Zinn-Justin S, Larrieu D. The BAF A12T mutation disrupts lamin A/C interaction, impairing robust repair of nuclear envelope ruptures in Nestor-Guillermo progeria syndrome cells. Nucleic Acids Res. 2022;50(16):9260–78.

71. Rzepecki R, Gruenbaum Y. Invertebrate models of lamin diseases. Nucleus. 2018;9(1):227–34.

72. Emond-Fraser V, Larouche M, Kubiniok P, Bonneil E, Li J, Bourouh M, et al. Identification of PP2A-B55 targets uncovers regulation of emerin during nuclear envelope reassembly in Drosophila. Open Biol. 2023;13(7):230104.

73. Torras-Llort M, Medina-Giro S, Escudero-Ferruz P, Lipinszki Z, Moreno-Moreno O, Karman Z, et al. A fraction of barrier-to-autointegration factor (BAF) associates with centromeres and controls mitosis progression. Commun Biol. 2020;3(1):454.

74. Duan T, Thyagarajan S, Amoiroglou A, Rogers GC, Geyer PK. Analysis of a rare progeria variant of Barrier-to-autointegration factor in Drosophila connects centromere function to tissue homeostasis. Cell Mol Life Sci. 2023;80(3):73.

75. Martin FA, Perez-Garijo A, Morata G. Apoptosis in Drosophila: compensatory proliferation and undead cells. Int J Dev Biol. 2009;53(8-10):1341–7.

76. Fan Y, Bergmann A. Apoptosis-induced compensatory proliferation. The Cell is dead. Long live the Cell! Trends Cell Biol. 2008;18(10):467–73.

77. Haynie JL, Bryant PJJWRsaodb. The effects of X-rays on the proliferation dynamics of cells in the imaginal wing disc of Drosophila melanogaster. 1977;183:85–100.

78. Wells BS, Yoshida E, Johnston LA. Compensatory proliferation in Drosophila imaginal discs requires Dronc-dependent p53 activity. Curr Biol. 2006;16(16):1606–15.

79. Kondo S, Senoo-Matsuda N, Hiromi Y, Miura M. DRONC coordinates cell death and compensatory proliferation. Mol Cell Biol. 2006;26(19):7258–68.

80. Garrido D, Bourouh M, Bonneil E, Thibault P, Swan A, Archambault V. Cyclin B3 activates the Anaphase-Promoting Complex/Cyclosome in meiosis and mitosis. PLoS Genet. 2020;16(11):e1009184.

